# Electrophysiological signatures of veridical head direction in humans

**DOI:** 10.1101/2023.01.26.525724

**Authors:** Benjamin J. Griffiths, Thomas Schreiner, Julia K. Schaefer, Christian Vollmar, Elisabeth Kaufmann, Stefanie Quach, Jan Remi, Soheyl Noachtar, Tobias Staudigl

**Author notes:** Corresponding Author: Tobias Staudigl.

## Abstract

Information about heading direction is critical for navigation as it provides the means to orient ourselves in space. However, given that veridical head direction signals require physical rotation of the head and most human neuroimaging experiments depend upon fixing the head in position, little is known about how the human brain is tuned to such heading signals. To address this, we asked fifty-two healthy participants undergoing simultaneous EEG and motion tracking recordings (split into two experiments) and ten patients undergoing simultaneous intracranial EEG and motion tracking recordings to complete a series of orientation tasks in which they made physical head rotations to target positions. We then used a series of forward encoding models and linear mixed-effects models to isolate electrophysiological activity that was specifically tuned to heading direction. We identified a robust posterior central signature that predicts changes in veridical head orientation after regressing out confounds including sensory input and muscular activity. Both source localisation and intracranial analysis implicated the medial temporal lobe as the origin of this effect. Subsequent analyses disentangled head direction signatures from signals relating to head rotation and those reflecting location-specific effects. Lastly, when directly comparing head direction and eye gaze-related tuning, we found that the brain maintains both codes while actively navigating, with stronger tuning to head direction in the medial temporal lobe. Together, these results reveal a taxonomy of population-level head direction signals within the human brain that is reminiscent of those reported in the single units of rodents.

## Main Text

Navigation is a complex cognitive phenomenon of which two basic forms can be distinguished: map-based and self-referential navigation (for review, see ^1^). While many have delved into how the human brain represents cognitive maps ^2–4^, comparatively few have explored how the human brain comprehends a sense of direction, a core ingredient to both forms of navigation. To redress this balance, we set out to explore how human electrophysiological activity is tuned to veridical (that is, true, physical) head direction.

Our understanding of the neural code for veridical head direction is based largely on rodent studies (though see ^5,6^), which demonstrate that single units across the brain are tuned to current heading direction ^7–12^. These “head direction” cells selectively fire when the rodent faces a particular angle in the environment, regardless of the physical location of the rodent in the environment ^13,14^, are persistent over time ^12,15^, continue in the absence of visual cues ^16^, and often precede physical head rotation ^17,18^. Furthermore, perturbation of the head direction cells destabilise representations of space, suggesting that representations of head direction are essential for navigation ^19–22^. Critically, given the abundance of head direction cells across the brain and the widespread, co-ordinated population activity they produce ^18,23,24^, representations of head direction should be detectable in macroscopic neuroimaging measures such as the local field potential (LFP; ^25–27^).

However, studying veridical head direction in humans is not a trivial task. Imaging methods such as magnetoencephalography (MEG) and magnetic resonance imaging (MRI) require the head to be fixed in position to minimise artefacts. Unfortunately, this prevents the generation of most self-referential cues, which is thought to limit the conclusions which can be drawn about active navigation ^28–30^. Electroencephalography (EEG), however, has no such limitation. For example, it has been used to successfully measure human brain activity during motor ^31,32^ and cognitive tasks ^33–35^ while the body is in motion. Here, we build on the success of these existing motion-based EEG studies to address how the human brain represents veridical head direction.

In the experiments presented here, we asked whether a signature of veridical head direction can be identified in the human scalp and intracranial EEG (iEEG). Participants completed a series of orientation tasks in which they made physical head rotations between multiple computer monitors, with each task containing a unique manipulation that helped disentangle head direction from other confounds (see figure 1a-b). We used forward encoding models to EEG activity as a function of simultaneously recorded heading angle, and then used linear mixed-effect models to localise the electrophysiological signals that were tuned to a given heading angle. We find converging evidence for a precisely tuned representation of head direction emanating from posterior regions including the parietal cortex and parahippocampus that is distinguishable from gaze-related neural codes and shares numerous similar properties to the head direction cells of rodents (as described above). In a second report, we investigate how these head direction-related signals are represented in memory and during sleep ^36^.

**Figure 1.**
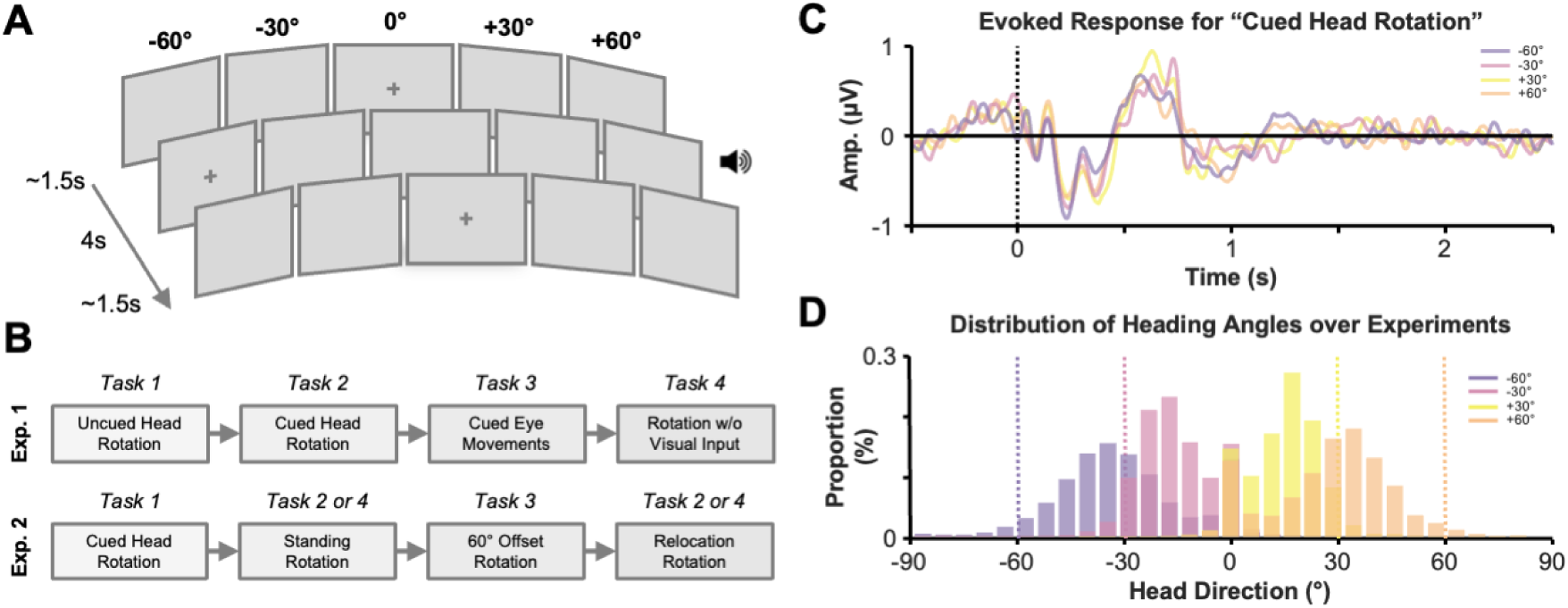
Experiment outline. **(A)** Participants completed a series of orientation tasks presented across five computer monitors. Participants began by fixating on a cross on the centre screen. This cross then moved to one of the other screens. In most tasks, participants were tasked with physically rotating their head towards the target screen. In most tasks, an auditory cue accompanied the movement of the fixation cross to help participants to track the cross. After a delay sufficient to allow participants to move into position and fixate, the cross returned to the centre screen. In experiment 2, an additional two screens were placed to the right of the five-monitor setup, with an angular orientation of 90° and 120° relative to the initial centre screen. **(B)** Three groups of participants completed four variations of these tasks. The first group of healthy participants and the group of patients completed Experiment 1. The second group of healthy participants completed Experiment 2. For full details of what each task entailed, see methods. **(C)** Visualisation of the evoked response at Pz following the onset of the head rotation in the “cued head rotation” task of Experiment 1. **(D)** Histogram of heading angles (relative to the centre screen) across all samples and participants of Experiment 1. Notable variability in heading direction can be seen here, meaning that the forward encoding models were able to make use of a wide range of heading angles rather than just 5 target conditions (i.e., -60°, -30°, 0°, +30° and +60°). Note that participants did not fully rotate their heads to the target screen; supplementary analysis instead suggests they made combined movements of the head and eyes to reach the target position (see supplementary figure 1).

## Results

### Scalp EEG activity is tuned to heading angle

In the first experiment, 32 participants (see table 1) completed four orientation tasks (see figure 1a-b), each with distinct manipulations used to delineate head direction-related activity from activity attributable to sensory cuing and visual input/eye movements (see supplementary figure 1 for head rotation time-series). We used cross-validated forward encoding models (FEM) to predict EEG activity based on head orientation. Head motion was continuously recorded and decomposed into basis sets of circular–Gaussian von-Mises distributions (from here on termed “kernels”) acting as model features in the FEM. Six of these FEMs were built, each using different kernel sizes (ranging from 6° to 60°) which allowed us to approximate the precision of the tuning of the EEG signal (see figure 1c) to current heading angle (see figure 1d). FEM performance was quantified by correlating the FEM-predicted EEG signal with the observed EEG signal (see supplementary figures 2-4 for topographies of these correlations). FEM performance was pooled across participants and linear mixed-effect models were used to assess how much variance current heading angle explained of the observed FEM performance after controlling for the co-occuring confounds of auditory input (i.e., animal sounds that pointed participants towards the target screens), visual input, eye movements, and muscular activity which the model would also be sensitive to (see methods for details).

For all tuning widths, EEG signal was tuned to changes in heading angle to a degree significantly greater than chance (peak z = 6.711, p_FDR_ < 0.001; see figure 2a-b). Descriptively speaking, FEM performance was maximal over posterior central sensors and when using a kernel width of 20° (see supplementary table 1 for statistics relating to all kernel widths and regressors; see supplementary figure 5 for statistical comparisons between kernel widths). When applying the FEM to source-space data, late visual and medial temporal regions responded most strongly to changes in heading angle (see figure 2c). While EEG activity was also tuned to muscular activity (peak z = 4.097, p_FDR_ < 0.001), the size of the effect was substantially smaller than that of head direction (see figure 2a-b). Importantly, the nature of the linear mixed-effect models means that this effect was statistically independent of that for head direction. Altogether, this suggests that activity present over the posterior central EEG is tuned to changes in heading angle – an effect that cannot be attributed to cuing, muscular activity or visual input/eye movements.

**Figure 2.**
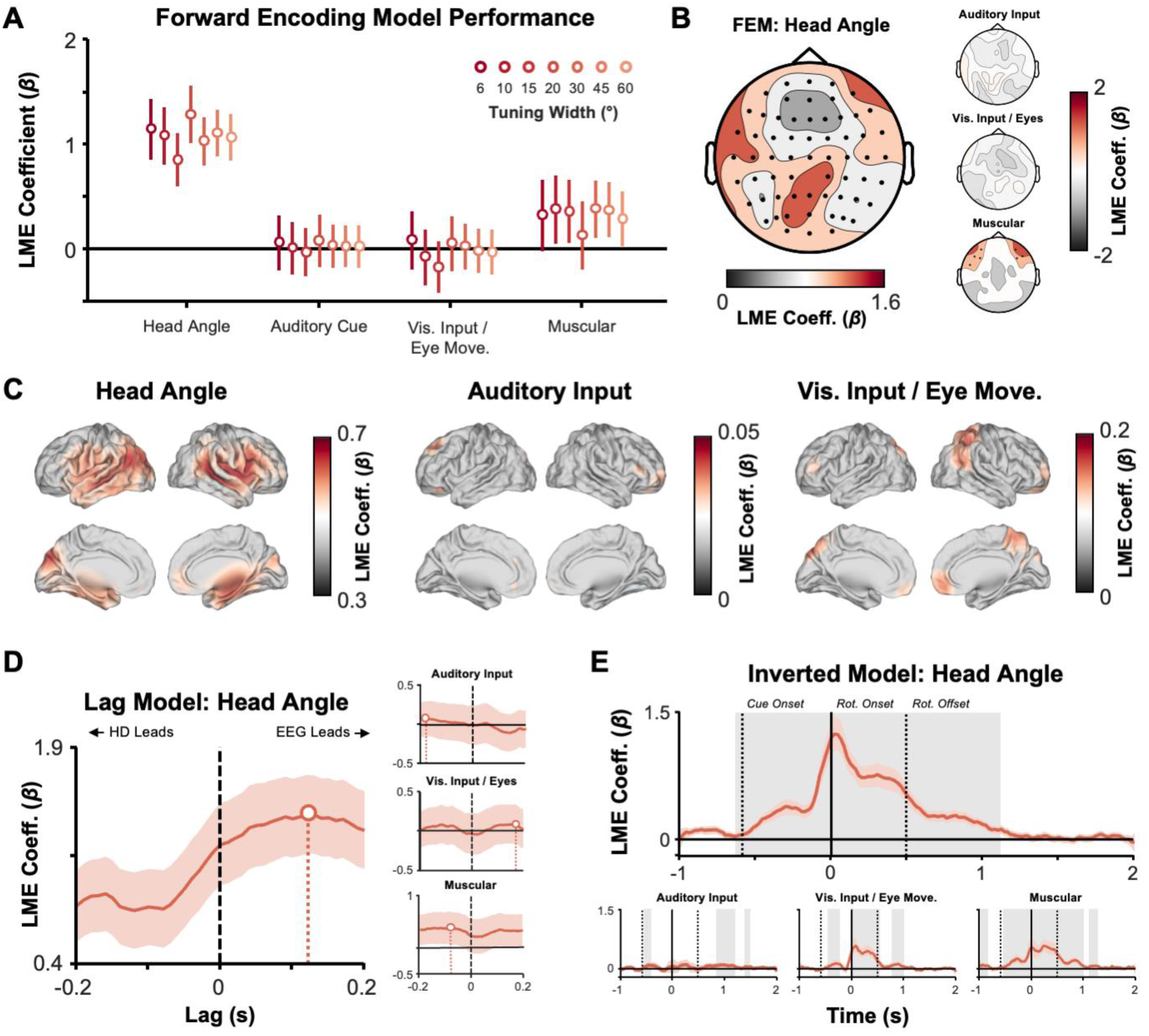
Electrophysiological activity tracks change in head angle. **(A)** Plot of the regressor coefficients (averaged over all electrodes) derived from linear mixed effects models used to predict forward encoding model performance. Dots indicate the estimated regressor coefficients. Error bars indicate the 95% confidence intervals of the coefficients, as computed by the Matlab function fitlme(). Colour indicates the tuning width of the kernel used in the forward encoding model, with darker colours representing the narrower tuning widths. For the head direction regressor, all tuning widths were significant (p_FDR_ < 0.05). **(B)** Topographic plots of the regressor coefficients derived from linear mixed effects models used to predict forward encoding model performance when using a 20° tuning width (i.e., the best performing forward encoding model). Deeper red colours indicate larger coefficient values. Black dots indicate electrodes that reliably map onto forward encoding model predictions (p_FDR_ < 0.05). Each of the four regressors are plotted separately. **(C)** Source plots of the regressor coefficients derived from linear mixed effects models used to predict forward encoding model performance when using a 20° tuning width. Deeper red colours indicate larger coefficient values. Each of the three regressors are plotted separately. Note that the EMG regressor was not included here as beamforming projects EMG components to outside the brain. **(D)** Time-series of regressor coefficients derived from linear mixed effects models used to predict lag-based forward encoding model performance. The deep red centre line indicates the estimated regressor coefficients. Error bars indicate the 95% confidence intervals of the coefficients around the estimated regressor coefficients. The dot indicates the lag at which predictive power of the regressor was greatest. “EEG Leads” refers to when changes in EEG activity precede changes in head direction; “HD Leads” refers to when changes in head direction precede changes in EEG activity..**(E)** Time-series of regressor coefficients derived from linear mixed effects models used to predict inverted encoding model performance. The deep red centre line indicates the estimated regressor coefficients. Error bars indicate the 95% confidence intervals of the coefficients around the estimated regressor coefficients. The shaded grey areas indicate significance (p_FDR_ < 0.05).

While these results demonstrate that EEG is tuned to heading angle, supplementary analyses found no evidence to suggest that individual EEG electrodes possessed a preferred direction tuning across participants, aligning with earlier work in rodents and human fMRI (e.g., ^6,27,37^; see supplementary figure 6)

While the previous analyses demonstrate that the patterns of neural activity that tune to changes in heading angle are statistically independent to those that tune to changes in visual input/eye movements, we wanted to complement these results by directly contrasting FEM performance between when the FEM was trained on heading angle data and when it was trained on gaze position data. In line with the results above, we found that neural activity tuned to both changes in heading angle and gaze position ^38,39^. Critically however, we found that posterior central neural activity was better predicted by changes in heading angle than by changes in gaze position (for full details, see supplementary figure 7).

Next, we examined the temporal dynamics of this tuned EEG activity, asking whether the EEG activity precedes or follows physical changes in heading angle. We first built a “lagged” encoding model, where the EEG time series was temporally shifted relative to the motion tracking time series prior to building the FEM. By repeating this for multiple temporal lags, we were able to build a time series which identified when the FEM best predicted EEG activity. Here, the lagged encoding model was best able to predict EEG activity when considering EEG activity that preceded heading angle information by approximately 120ms (peak z = 8.371, p_FDR_ < 0.001; see figure 2d). A direct contrast of FEM performance for timepoints when EEG led or lagged heading angle supported this observation, with the encoding model performing significantly better for samples when EEG activity preceded the change in heading angle relative to when it followed the change (z = 9.83, p_FDR_ < 0.001). This aligns with animal and computational work suggesting that signatures of head direction are anticipatory rather than responsive ^17,18^.

We complemented our lagged FEM with an inverted encoding model, which takes the weights from the training set and inverts them to produce an estimate of heading angle for every sample. Representational similarity analysis ^40^ was then used to see when the predicted head angle differed most greatly between the four outside monitor positions. In line with the lagged FEM, the inverted encoding model showed that the predictive power of EEG activity for heading angle ramped up just before the head began to rotate (peak z = 9.480, p_FDR_ < 0.001; see figure 2e). EEG activity continued to predict heading angle throughout the head rotation and beyond as participants maintained the new heading angle (albeit to a slightly lesser degree). This aligns with animal work showing persistent but weaker head direction tuning after head rotation ^12^. Importantly, persistent coding in the absence of the physical movement rules out the possibility that the effects reported at the beginning of this section are a result of angular head velocity rather than heading angle (for further details, see supplementary figure 8). Together, these results suggest that tuned EEG activity precedes the change in veridical heading angle and is sustained after the head rotation is completed.

Notably, the anticipatory EEG effects are not simply a manifestation of associative links between the auditory cue and particular head rotations. In an isolated analysis of the “uncued head rotation” task, which occurred prior to any association being formed between the cues and the head rotations, the lagged encoding model continued to best predict EEG activity when EEG activity preceded changes in heading angle (max. z = 5.934, p_FDR_ < 0.001; lead vs. lag contrast: max. z = 5.183, p_FDR_ < 0.001; see supplementary figure 9). This suggests that the anticipatory nature of the EEG tuning-related activity observed here is not solely attributable to associative memory.

### Intracranial EEG activity is tuned to heading angle

Several key regions expressing head direction cells are found deep within the brain, which can be difficult to reliably measure in scalp EEG recordings. Therefore, we set out to repeat the first experiment in a group of ten patients with intracranial depth electrodes. The methodological and analytical approaches matched that of the scalp EEG experiment, with the single exception of the linear mixed-effect models. As patients have bespoke implantation schemes, models cannot be built for individual electrodes as done above. Therefore, we built models for several regions-of-interest (ROI) instead; one ROI for each cortical lobe of the brain, and one for each recorded subregion of the medial temporal lobe (a decision driven by the fact that subregions of the medial temporal lobe have unique functions in navigation^2^ [see methods for details]). As before, individual patients were modelled as random effects.

When probing individual lobes, all regions of interest demonstrated a tuned response to current heading angle except the occipital lobe (frontal lobe: peak tuning width = 60°, z = 5.508, p_FDR_ < 0.001; parietal lobe: peak tuning width = 6°, z = 2.844, p_FDR_ < 0.001; occipital lobe: peak tuning width = 60°, z = 1.721, p_FDR_ = 0.110; lateral temporal lobe: peak tuning width = 10°, z = 5.306, p_FDR_ < 0.001; see figure 3a; see supplementary figure 10 for correlations between iEEG activity and FEM predictions; see supplementary figure 11 for tuning width comparisons). In the case of the occipital lobe, this region instead appeared to show substantial tuning to saccadic activity/visual input (peak tuning width: 15°, z = 2.704, p_FDR_ = 0.015). Descriptively speaking, the lag-based models of the intracranial recordings peaked such that changes in iEEG signal preceded changes in head angle for the parietal and temporal lobes (parietal lobe: peak lag = 190ms, z = 4.832, p_FDR_ < 0.001; lateral temporal lobe: peak lag = 120ms, z = 5.554, p_FDR_ < 0.001; see figure 3c). In the frontal lobe, changes in head angle preceded the iEEG signal (peak lag = - 100ms, z = 6.145, p_FDR_ < 0.001). Furthermore, the parietal and temporal models performed significantly better when iEEG activity preceded the change in heading angle relative to when it followed the change (parietal lobe: z = 2.052, p = 0.015; lateral temporal lobe: z = 2.230, p = 0.025). No similar directional effect was observed in the frontal lobe (frontal lobe: z = 0.490, p > 0.5).

**Figure 3.**
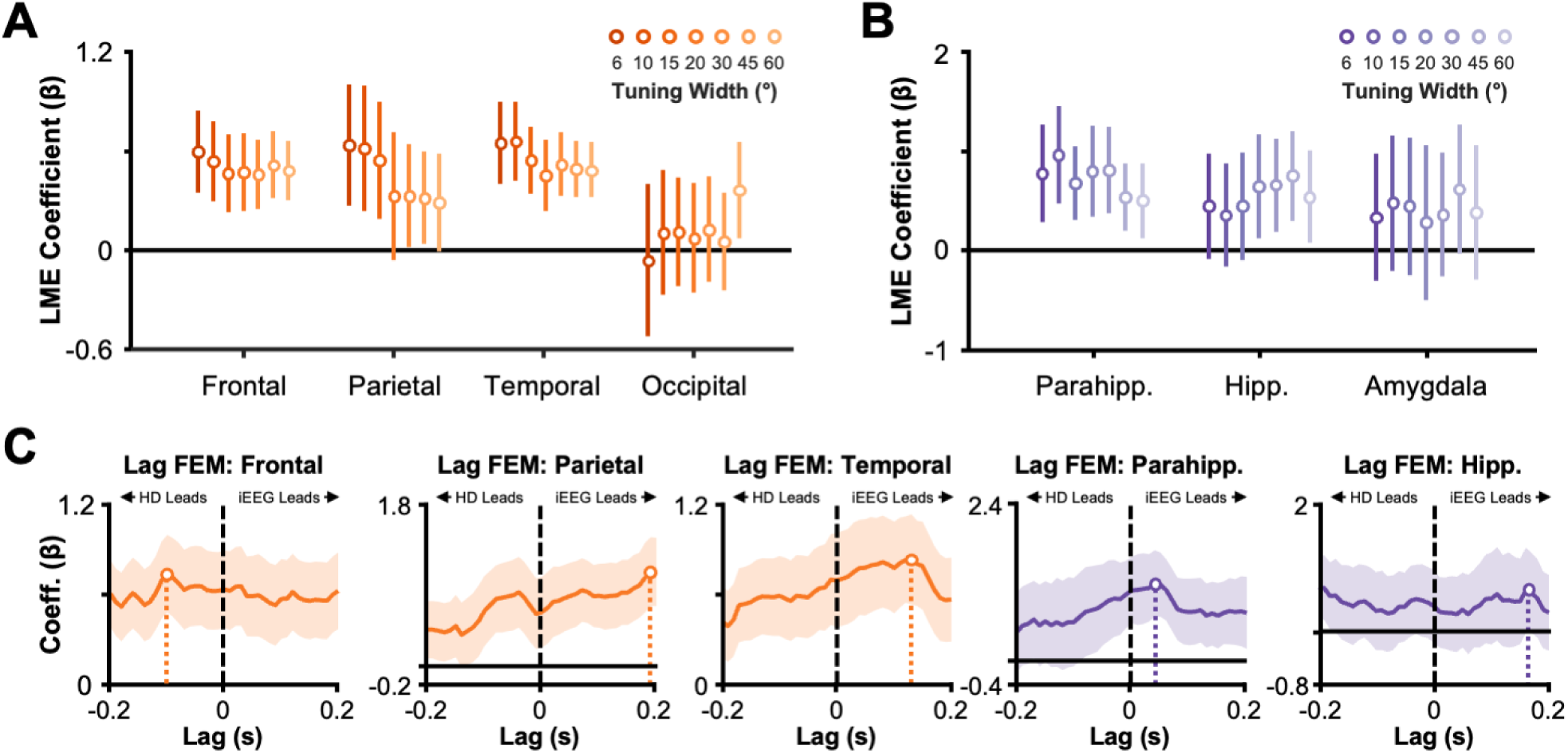
Intracranial electrophysiological activity tracks change in heading angle. **(A)** Plot of the head direction regressor coefficient derived from linear mixed effects models used to predict forward encoding model performance. Dots indicate the estimated regressor coefficients. Error bars indicate the 95% confidence intervals of the coefficients around the estimated regressor coefficients, as computed by the Matlab function fitlme(). Colour indicates the tuning width of the kernel used in the forward encoding model, with darker colours representing the narrower tuning widths. For the frontal, parietal and temporal lobes, all tuning widths were significant (p_FDR_ < 0.05). **(B)** Details of plot match that of panel A, but here plotted for subregions of the medial temporal lobe. Parahippocampus and hippocampus demonstrated significant predictive power for tracking changes in head orientation (parahippocampus: p_FDR_ = 0.035; hippocampus: p_FDR_ = 0.015;). **(C)** Time-series of regressor coefficients derived from linear mixed effects models used to predict lag-based forward encoding model performance. Plots are restricted to those regions which demonstrated significant predictive power in the previous analysis. The deep coloured centre line indicates the estimated regressor coefficients. Error bars indicate the 95% confidence intervals of the coefficients, as computed by the Matlab function fitlme(). The dot indicates the lag at which predictive power of the regressor was greatest.

When probing the subregions of the medial temporal lobe, both the hippocampus and parahippocampus demonstrated substantial tuning to current head angle (parahippocampus: peak tuning width = 30°, z = 2.517, p_FDR_ = 0.035; hippocampus: peak tuning width = 45°, z = 2.752, p_FDR_ = 0.015; amygdala: peak tuning width = 10°, z = 1.105, p_FDR_ = 0.298; see figure 3b). As with the lobe-level analysis, the lag-based models of the iEEG signal suggested that changes in both the hippocampal and parahippocampal signal preceded changes in head angle (hippocampus: peak lag = 160ms, z = 2.657, p_FDR_ < 0.001; parahippocampus: peak lag = 40ms, z = 3.637, p_FDR_ < 0.001; see figure 3c). However, in both cases, the peaks were less clear than in the lobe-based or scalp-level analyses, and this is reflected in a direct contrast of model performance, which found no significant difference in model performance between when iEEG led or lagged heading angle (parahippocampus: z = 0.852, p = 0.185; hippocampus: z = 0.086, p > 0.5).

Taken together, these results suggest that several distinct regions are tuned to changes in head angle, including the parietal lobe and parahippocampus. While the lag-based effects were less clear-cut than their scalp EEG equivalents, the parietal and temporal lobes continued to show a significant effect where iEEG activity preceded changes in heading angle and the remaining regions continued to showed a descriptive trend in which model performance peaked when iEEG activity preceded the change in heading angle. Notably, within the medial temporal lobe, parahippocampus but not hippocampus showed substantial directional tuning, which aligns with animal work demonstrating HD cells outside hippocampus ^41^.

### Conceptual replication of heading angle-tuned EEG activity

In the second experiment, 20 participants completed four orientation tasks that varied the position of the participants relative to the monitors. Two of these tasks involved sitting while the others involved standing (introducing a position change in the z-axis). One of the standing tasks also involved standing one metre away from the standard location (introducing a location change in the x-axis), while one of the sitting tasks involved sitting in the same location but rotated by 60° (a condition that will be discussed in more detail in the last results subsection). In the first instance, we used the same approach to forward encoding modelling as in Experiment 1 to both replicate and extend those findings. Once again, linear mixed-effect models were used to delineate the influence of head angle from potential confounds, in this case: position in the z-axis, location in the x-axis, and muscular activity.

Mapping onto the findings from Experiment 1, we found that, after regressing out the influence of location and muscular activity, scalp EEG recordings demonstrated a tuning to current head angle significantly beyond what would be expected by chance (peak z = 7.063, p_FDR_ < 0.001; see figure 4a-b; see supplementary table 2 for statistics relating to all kernel widths and regressors; see supplementary figure 12 for tuning width comparisons). These effects were most prominent over posterior central electrodes and were localised to occipital and medial temporal regions (albeit not reaching as far anteriorly as in Experiment 1; see figure 4c). Both the lag-based forward encoding model and the inverted encoding model also produced results matching Experiment 1, suggesting that changes in EEG activity precede changes in head angle (lag model: peak z = 5.37, p_FDR_ < 0.001; lead/lag contrast: z = 6.58; p_FDR_ < 0.001; inverted model: peak z = 6.13, p_FDR_ < 0.001; see figure 4d-e).

**Figure 4.**
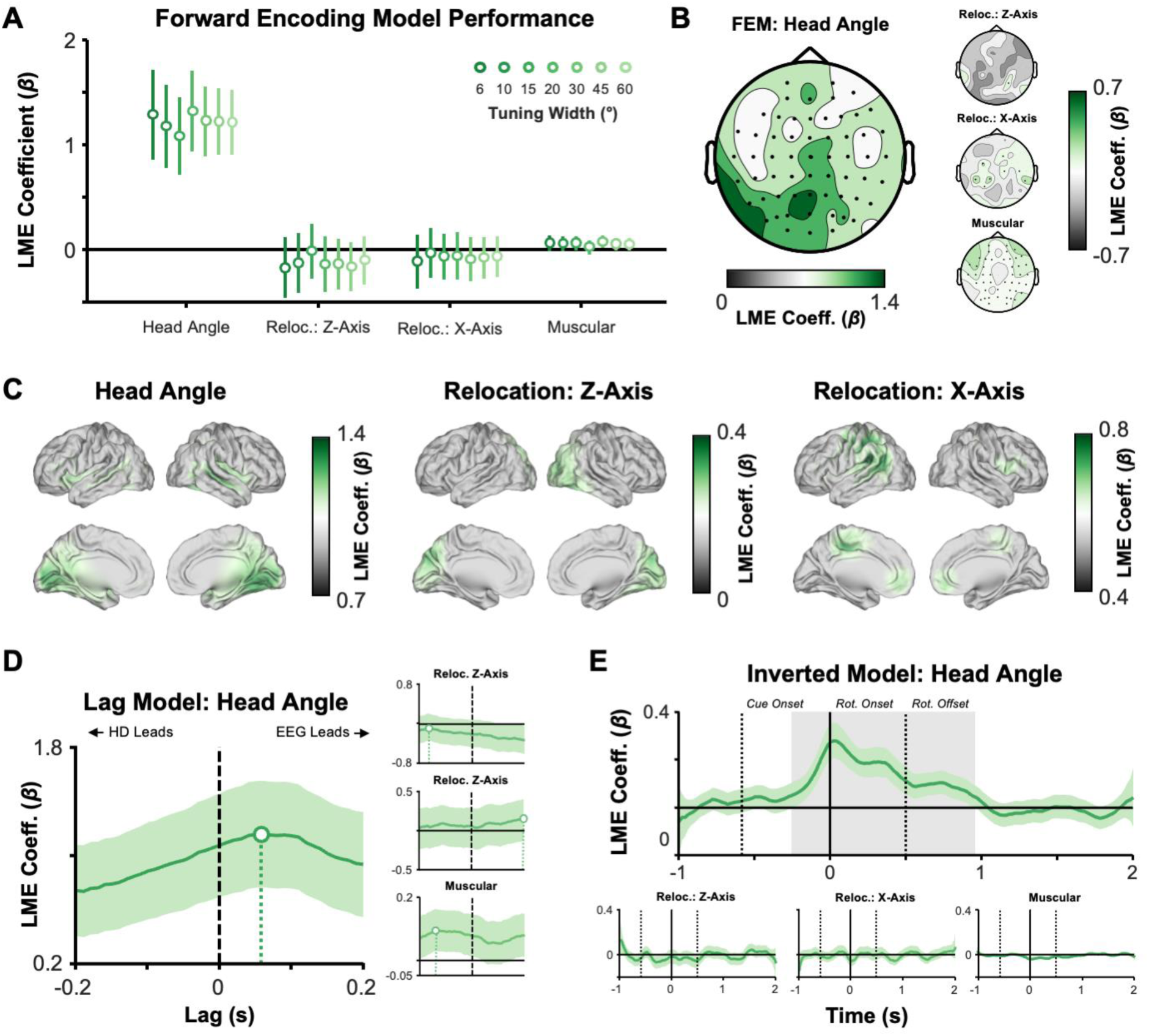
Electrophysiological activity tracks change in head angle independent of location. **(A)** Plot of the regressor coefficients derived from linear mixed effects models used to predict forward encoding model performance. Dots indicate the estimated regressor coefficients. Error bars indicate the 95% confidence intervals of the coefficients around the estimated regressor coefficients, as computed by the Matlab function fitlme(). Colour indicates the width of the kernel used in the forward encoding model, with darker colours representing the narrower tuning widths. For the head direction regressor, all tuning widths were significant (p_FDR_ < 0.05). No other regressor produced significant effects. **(B)** Topographic plots of the regressor coefficients derived from linear mixed effects models used to predict forward encoding model performance when using a 20° tuning width (i.e., the best performing forward encoding model). Deeper green colours indicate larger coefficient values. Black dots indicate electrodes that reliably map onto forward encoding model predictions (p_FDR_ < 0.05). Each of the four regressors are plotted separately. **(C)** Source plots of the regressor coefficients derived from linear mixed effects models used to predict forward encoding model performance when using a 20° tuning width. Deeper green colours indicate larger coefficient values. Each of the three regressors are plotted separately. Note that the EMG regressor was not included here as the process of beamforming will inherently project EMG components to outside the brain. **(D)** Time-series of regressor coefficients derived from linear mixed effects models used to predict lag-based forward encoding model performance. The deep green centre line indicates the estimated regressor coefficients. Error bars indicate the 95% confidence intervals of the coefficients around the estimated regressor coefficients, as computed by the Matlab function fitlme(). The dot indicates the lag at which predictive power of the regressor was greatest. **(E)** Time-series of regressor coefficients derived from linear mixed effects models used to predict inverted encoding model performance. The deep green centre line indicates the estimated regressor coefficients. Error bars indicate the 95% confidence intervals of the coefficients around the estimated regressor coefficients, as computed by the Matlab function fitlme().. The shaded grey areas indicate significance (p_FDR_ < 0.05).

The findings of Experiment 1 and their replication reported above suggest that human EEG activity is tuned to changes in current heading angle. However, it would be premature to suggest that, based on these results alone, this reflects a human analogue of the prototypical head direction signal observed in rodents. Specifically, we must probe whether the head direction effects uncovered here are (1) consistent over changes in location, and (2) separable from effects attributable to head rotation. The next two subsections address these issues in turn.

### Tuned EEG activity persists across locations

To delineate the effects of location-independent and location-specific head direction signals, we adapted the forward encoding modelling approach to consider how the models generalised across tasks. This generalisation was achieved by training the FEM on one task and applying it to data from another task (for a similar example of this approach, see ^42^). For EEG electrodes where the FEMs do generalise to tasks completed in differing locations (that is, the FEM performs better than what would be expected by chance), one could conclude that the EEG signal predicted by the FEM is not dependent on location. For electrodes where the models perform well within a given task but do not generalise, one could conclude that the head angle effect is specific to a given location. Linear mixed-effects models were used to explore the influence of location on head direction signal, as well as regress out the influence of muscular activity (as done in the previous analyses).

This approach revealed that the generalised FEMs could reliably predict posterior central EEG activity (peak z = 4.540, p_FDR_ < 0.001; see figure 5a), suggesting they make use of location-independent heading angle information. Analysis of source-space data suggested these effects originated from right visual ventral, right parahippocampal and left inferior frontal areas (see figure 5b). Notably, the linear mixed-effects models also found evidence to suggest some location-specific head direction-related signals exist within the EEG (peak z = 7.062, p_FDR_ < 0.001). These effects were positioned slightly further back on the EEG cap, with source analysis implicating the posterior parietal cortex and occipital lobe (see figure 5a-b). These results suggest that distinct parts of the EEG signal are tuned to location-independent and location-specific signatures of current heading angle.

**Figure 5.**
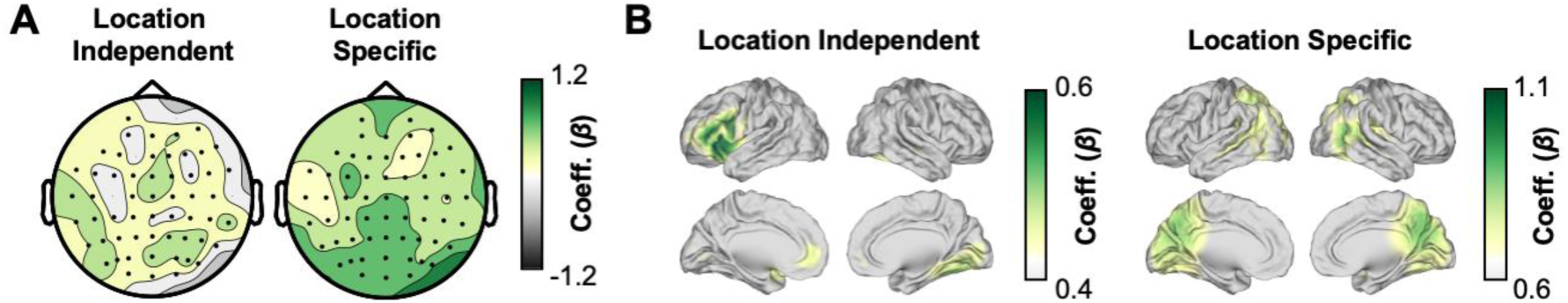
Electrophysiological representations of location-independent and location-dependent head direction signals. **(A)** Topographic plots of the two regressor coefficients derived from linear mixed effects models used to predict forward encoding model performance when using a 20° tuning width (i.e., the best performing forward encoding model). Deeper green colours indicate larger coefficient values. Black dots indicate electrodes that reliably map onto forward encoding model predictions (p_FDR_ < 0.05). **(B)** Source plots of the two regressor coefficients derived from linear mixed effects models used to predict forward encoding model performance when using a 20° tuning width. Deeper green colours indicate larger coefficient values.

### Tuned EEG activity is distinct from rotation-related activity

A similar approach was used to delineate EEG activity tuned to head direction (i.e., that anchored to the environment) from EEG activity tuned to head rotation (i.e., that anchored to the body). In this instance, FEMs from one sitting condition were generalised to the other sitting condition (where participants were rotated 60° to the right in the environment), and vice versa. If the FEMs generalise, it would indicate that the effect is driven by head rotation (as the model would be insensitive to the 60° rotation of the entire body within the environment). In contrast, any part of the FEM that does not generalise would reflect head direction. Here, the linear mixed-effects models suggested that the head direction effects in previous analyses were a combination of “true” head direction signals and those tied to the rotation of the head. The “true” head direction signals were the most prominent over outer occipital and temporal electrodes (peak z = 4.679, p_FDR_ < 0.001; see figure 6a), with source-space analyses implicating the visual ventral stream and temporal pole (see figure 6b). The head rotation signals were most prominent over posterior central electrodes (peak z = 4.92, p_FDR_ < 0.001), and were localised to the posterior parietal cortex and frontal lobe (see figure 6a-b). These results suggest that both “true” head direction signals and signals relating to head rotation are present in EEG activity.

**Figure 6.**
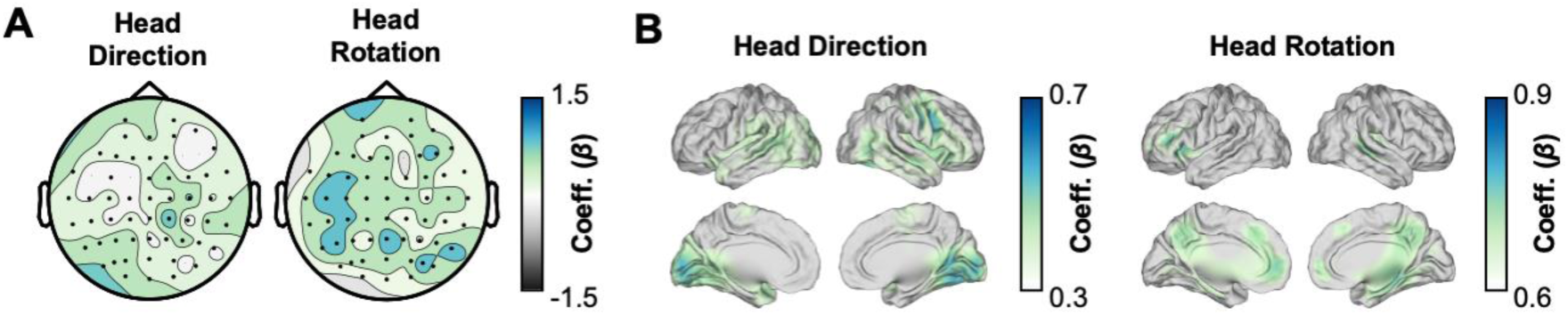
Electrophysiological representations of environment-based head direction and body-based head rotation signals. **(A)** Topographic plots of the two regressor coefficients derived from linear mixed effects models used to predict forward encoding model performance when using a 20° tuning width (i.e., the best performing forward encoding model). Deeper blue colours indicate larger coefficient values. Black dots indicate electrodes that reliably map onto forward encoding model predictions (p_FDR_ < 0.05). **(B)** Source plots of the two regressor coefficients derived from linear mixed effects models used to predict forward encoding model performance when using a 20° tuning width. Deeper blue colours indicate larger coefficient values.

## Discussion

To date, it has remained unclear how the human brain is tuned to changes in veridical head direction, and where such tuning would be localised. To remedy this, we asked participants to complete a series of head rotation tasks while undergoing scalp or intracranial EEG recordings with the aim of isolating the neural signatures of human veridical head direction. Using a series of analytical models (see figure 7), we identified a signature tuned to current heading angle that is distinguishable from sensory input and muscular activity. Anatomically speaking, both source-localised scalp EEG and intracranial EEG implicated a wide network of regions in this tuning, including the medial temporal lobe and parietal cortex. These findings were replicated in a second, independent sample of healthy participants, who completed a series of additional tasks that regressed out the influence of head rotation and location-specific effects to further isolate a neural signature specifically related to veridical head direction. Altogether, these results provide a detailed taxonomy of head direction-related signals that can be observed in free-moving human participants.

**Figure 7.**
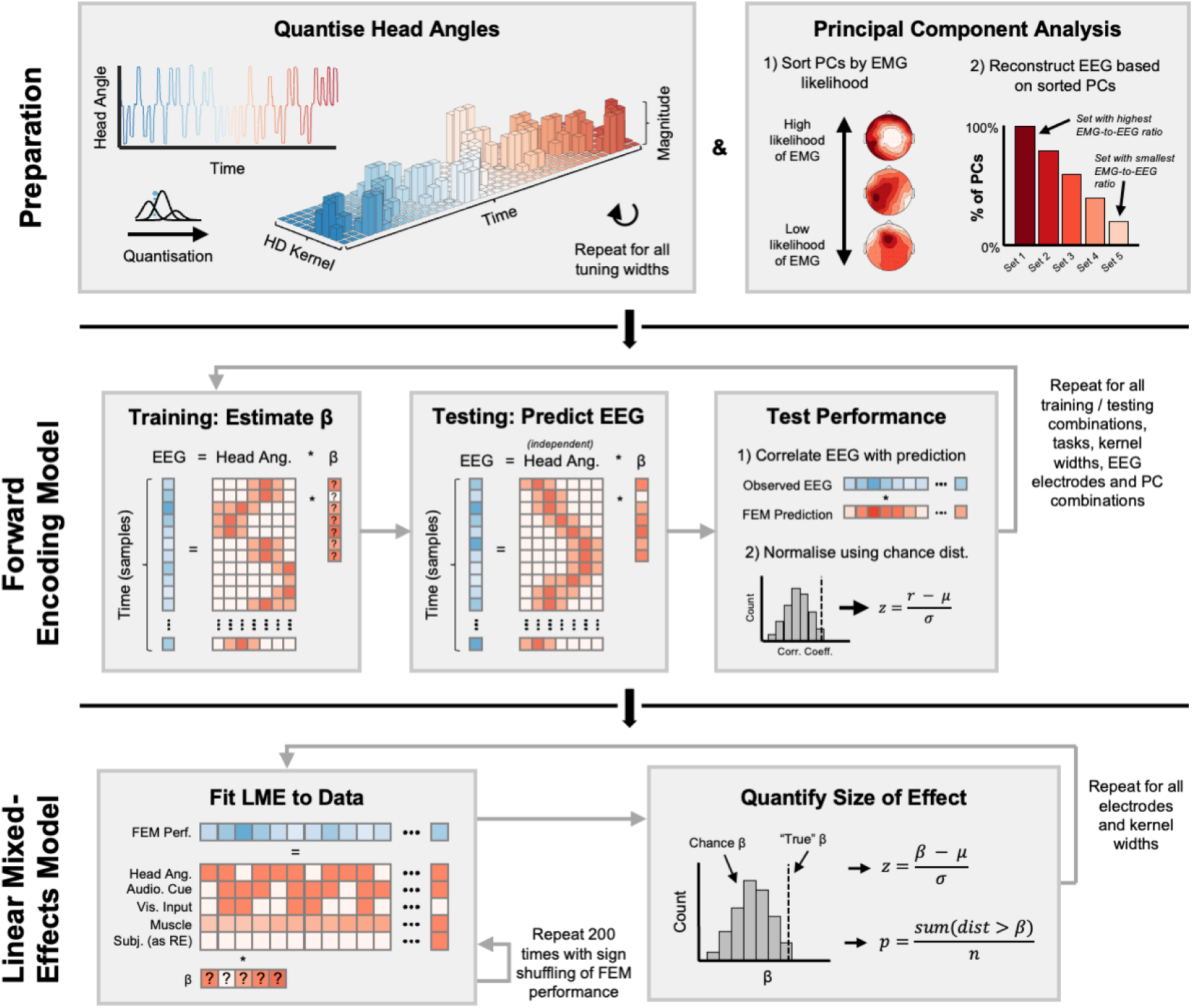
Visual depiction of analysis pipeline for Experiment 1. The analysis began with the preparation of motion tracking and EEG signals. The continuous, univariate motion tracking signal was convolved with a series of kernels to provide a 2-dimensional design matrix where each column reflected a single kernel and each row reflected a single point in time (see top left). EEG was broken down into principal components and then reconstructed using one of five proportions of components that were least likely to reflect electromuscular (EMG) activity (see top right). The forward encoding model (FEM) was then trained, tested and validated (see middle row). The motion and EEG data were split into four folds. The model was trained on three of these folds (see middle left). The resulting model weights were then applied to the held-out fold and used to estimate EEG activity for this fold (see centre of middle row). The model was then validated by correlating the predicted EEG activity if the real EEG activity of the held-out fold (see middle right). The resulting correlation co-efficient was normalised against a chance distribution of correlation values derived from randomly time-shifting the predicted EEG time series relative to the real EEG activity. Lastly, linear mixed-effects (LME) models were used to investigate how distinct latent factors influenced FEM performance (see bottom row). The LME models were fit to the observed (or “true”) data and to 200 instances of data were the sign of the FEM performance was flipped at random (see bottom left). The size of the resulting effect was then computed by comparing the LME weights derived from the “true” data to the chance data (see bottom right).

### On the (dis)similarities to rodent head direction cells

For almost four decades, animal research has steadily advanced our knowledge of the electrophysiological signatures of veridical head direction signals ^e.g., 6,11,43^. Here, we aimed to expand this work to electrophysiological population activity in humans and found numerous similarities (and the occasional idiosyncrasy) to what has been reported in animal models. Below, we compare our results from human population-level activity against what could be considered some of the main features of rodent head direction cells:

1. *Anatomical overlap*: We observed directional tuning to current head angle in many regions across the cortex. The strongest of these effects arose in the medial temporal lobe, where head direction cells are observed in abundance^6,11,43^. Importantly, source-localization of scalp EEG converged with intracranial EEG recorded directly in the medial temporal lobe, providing coherent evidence from two different recording techniques in two independent cohorts. Zooming in on the subregions of the medial temporal lobe that were recorded intracranially, we observed directional tuning outside but not in the hippocampus, which is in line with many animal studies^41^. Since the implantation of iEEG electrodes is restricted by clinical necessity, we were not able to record from other brain regions that, in the rodent, host head direction cells in abundance (e.g., the anterior thalamus^12^; note that scalp EEG is unlikely to spatially resolve signals from such a small and deep structure). We did, however, observe tuning effects in the frontal lobe, a region not featured in many maps of the head direction network^11^. During navigation, activity in the frontal lobe has been attributed to several functions, such as goal tracking and route planning ^44^. Given that our experiment involved tracking the target direction and intentionally orienting towards the target, the heightened tuning of frontal activity to current heading angle (relative to tuning during free exploration in animal studies) is perhaps not surprising. In sum, while we did observe some regional selectivity, our results more generally align point towards head direction activity being observable across the brain, aligning with numerous reports indicating that head direction-related activity can be observed in many regions across the brain^6,11,43^.
2. *Independence from visual input*: Directional tuning of head direction cells is maintained when visual input is removed ^16^. In our experiment, using virtual reality goggles containing nothing but a fixation cross (eliminating changes in visual input that could be used as visual cues as well as minimizing eye movements) and linear mixed-effect modelling, we could separate the influence of visual input and eye movements from that of head direction-related activity. While visual input-related activity was centred on superior occipital and inferior parietal regions, activity tuned to current head direction was centred in the medial temporal lobes. The fact that population-level directional tuning was sustained in the absence of visual input brings the current results further in line with the traditional definition of head direction-related activity ^16^.
3. *Independence from location*: Cells tuned to head direction can be classified as “traditional” head direction cells that fire independently of the animal’s location within an environment or “sensory” cells that are dependent on sensory input ^43^. We observed similar dissociable patterns of activity that relate to these two signatures of head direction: “traditional” location-independent head direction-related activity stemmed from the visual ventral regions, the parahippocampus, and left inferior frontal gyrus, while location-specific head direction-related activity was more closely related to activity in the posterior parietal cortex. Notably, this spatial delineation overlaps with work in rodents, which identified location-sensitive head direction-related activity within the retrosplenial cortex and the anteroventral thalamus ^45,46^. Evidence that population-level directional tuning was sustained across locations further supports the idea that a portion of the tuned activity we observed matches that of “traditional” head direction activity.
4. *Anchoring to the environment*: Head direction cells are anchored to the position of the head within the environment, not to the position of the head with respect to the body ^13^. We separated “body-based” tuning to head rotation from more traditional head direction effect through experimental and statistical means and identified distinct hubs of activity for the two processes. Once again, head direction-related activity was most prevalent in the visual ventral and temporal regions. In contrast, “body-based” rotation-related activity was observed in the posterior parietal cortex (see figure 6b), aligning with reports that posterior parietal lesions do not impact tuning to head direction ^47^ and supporting the idea that the posterior parietal cortex is instead involved in translational updating ^48^.
5. *Anticipatory signatures*: In line with animal research, we observed the tendency for tuned signatures of head direction to precede the physical head rotation ^49,50^. Specifically, in both scalp EEG datasets, model performance was significantly better when EEG led rather than lagged changes in head direction (though this effect was less clear in the intracranial recordings, perhaps due to reduced statistical power available in the smaller sample sizes). Intriguingly, the delay between neural signature and physical head rotation observed here most closely matched the delay exhibited by the lateral mammillary nucleus (∼40-100ms; positioned at the early stages of the head direction network) and not the delay exhibited by the postsubiculum (located later in the head direction network hierarchy, showing almost instantaneous responses to changes in head direction ^17^), despite the regions identified in our task being situated closer to the postsubiculum than the lateral mammillary nucleus. It is unclear why this is the case, but perhaps (as with the frontal lobe effects discussed above) the fact that participants were cued to rotate to a particular target meant that anticipatory responses are earlier than those observed during free exploration.
6. *Tuning widths*: In rodents, individual head direction cells show a directional tuning of around 60 to 120° ^13,14^. However, the population-level tuning observed here was, descriptively speaking, more precise (∼10-20°; aligning with similar reports from VR studies of human directional tuning; ^27^. This discrepancy may relate to the fact that tuning width tends to become more precise as one progresses further down the head direction network hierarchy ^50,51^. Given that scalp EEG will principally pick up on cortical activity at the bottom end of this hierarchy, it is perhaps no surprise that the tuning we observed is more precise. This idea is supported by descriptive interpretation of the intracranial data, which suggests that the medial temporal lobe is more coarsely tuned (∼15°) than neocortical sources (e.g., ∼10° tuning in parietal lobes), as well as 7T fMRI work suggesting a gradient of tuning widths along the visual ventral stream which becomes coarser as one approaches the medial temporal lobe ^27^.
7. *Persistence in the absence of movement*: In rodents, head direction cells fire persistently for at least ∼600ms after head rotation has ceased ^15^. Our inverted encoding models came to similar conclusions, suggesting that tuned population activity is sustained for ∼500ms after the offset of the head rotation (see figure 2e). Notably, however, tuned activity dissipates shortly after this. While not completely contradictory to the observed decline in head direction cell firing rates that accompany immobility ^52^, the result is hard to reconcile with influential computational models which suggest that head direction-related activity should be persistent in the absence of movement (for review, see ^12^). We speculate this decline in tuning relates to participants anticipating the return to the centre screen, which would produce a signature of the imagined upcoming rotation ^53^, impairing the inverted encoding model’s ability to predict current head angle.
8. *Population-level activity*: The most egregious dissimilarity between our results and those reported in rodents is the difference between population-level tuning and single-cell tuning. To what extent is tuned population-level activity reported here comparable to single-cell tuning? We speculate that there is a close relationship, with population-level activity arising as an emergent property of a network of head direction cells synchronising their action potentials. Such population-level activity may be essential for propagating head direction-related information across the head direction network hierarchy as synchronous firing would ensure neurons effectively impact downstream regions in the hierarchy ^6^. Whether the level of synchronicity between head direction cells is relevant for behaviour, however, remains an intriguing open question.

Taken together, population-level activity tuned to current heading angle in free-moving humans shares several similar properties with individual head direction cells in animal models. However, open questions remain. Some pertain to the discrepancies between what was observed here and what has been reported in previous studies (see above). Others relate to facets of head direction cells not explored here. For example, head direction cells continue to fire when rodents are passively moved through an environment ^54^, but it remains unknown if the same is true for population-level activity in humans. Moreover, the goal-directed nature of our task differs from free exploration used in rodent studies (and may explain differences related to anticipatory timing and persistence). It would be of interest to compare human population-level tuning to rodent single-unit tuning in tasks involving free exploration. Nonetheless, we feel that the similarities outweigh the differences, leading us to suggest that we can measure an analogue of head direction in population-level activity of free-moving humans.

### On related human/non-human primate head direction studies

Compared to the vast literature on head direction cells in rodents, few studies in non-human primates ^39,55,56^ and, to the best of our knowledge, no study in humans has reported tuning of neural activity to veridical head direction. Instead, a growing number of studies hint that human and non-human primates are more visually-orientated when navigating, and may utilise a grid-like code to navigate visual space ^38,57–64^. Indeed, there are reports that, within the macaque entorhinal cortex, a subset of cells are tuned to the direction of saccades ^65^. Our results do not refute these claims, but do suggest that rodent-like head direction signals co-exist with these visual/saccadic representations of space, in line with work in non-human primates showing a mix of head- and eye-gaze coding ^38,39^ with potentially overlapping circuitry involved in coordinating head and eye movements ^66–70^. Using linear mixed effects models, we simultaneously modelled (and, consequently, delineated) the influence of head direction and saccade direction on neural tuning, revealing that the medial temporal lobe is intimately tuned to head direction (see figure 2C). While it remains an open question as to (i) whether head- or saccade-direction tuning takes precedence during free navigation, and/or (ii) the brain can flexibly switch between the two codes, our results suggest that the human brain maintains head direction-codes while actively navigating.

Our observation of head-direction tuned neural activity in the free-moving human brain aligns with several previous studies that have utilised virtual reality in conjunction with methodologies such as fMRI and single unit recordings. ^71–75^. Of particular relevance here, a recent fMRI study used forward encoding models to model “virtual head direction” information (that is, heading information within a virtual environment) and found evidence for the representation of current heading direction within ventral occipital, medial parietal and medial temporal areas ^27^. However, as participants were immobilised by necessity, the critical contributions of vestibular input to head direction information ^52,76,77^ were excluded, questioning whether a full neural signature of head direction could be observed ^28–30^. Despite this conceptual concern, the fMRI findings align with our results, implicating regions such as the parahippocampus in the representation of head direction.

However, some inconsistencies between our work and past research do exist, and may be attributable to task differences. For example, in many virtual-reality-based studies, participants are required to self-navigate through a virtual environment principally relying on vision (a necessity due to the immobilisation required to record the neural data). In contrast, our free-moving task allowed participants to utilise non-visual sensory cues and, in some instances, explicitly prevented the use of visual sensory information. The comparatively heavy emphasis on vision that is incurred in virtual-reality studies may explain why Nau and colleagues (2020) observed their strongest tuning in early occipital regions. In a similar vein, the dispersed head-direction related signals we observed in free-moving participants may be attributable to the comparatively wide range of sensory cues available leading to many more regions becoming involved in the task at hand.

Of course, these past studies hold an advantage over our study in that they sample a full range of head directions (i.e., -180° to +180°) whereas we have only sampled a subset of these angles (-60° to +60° in Experiment 1; -60° to +120° in Experiment 2). However, as there is no evidence to suggest that the brain uses distinct mechanisms to code for distinct head directions, and it is hard to argue why the brain would indeed evolve to do so, we do not consider it a major issue that we only sample a subset of head directions here and would speculate that our results generalise to a full range of head directions.

A growing number of studies are incorporating full body movements with EEG recordings (also known as “mobile brain/body imaging” ^78^ or “mobile cognition” ^28^) to assess how brains track naturalistic movement. Importantly, several studies have explored how brain activity related to head rotation differs from other forms of body movement or when the head is stationary ^31,32^. While these studies implicated the retrosplenial cortex in tracking changes in heading direction, the nature of these experiments prohibited detailed conclusions about whether retrosplenial activity was tuned to particular heading directions in the environment, was tuned to patterns of gaze activity, or was agnostic to both and instead coded for changes in heading rotation. By combining the analytical approaches previously used in fMRI ^27,79^ with the methodological approaches used in mobile brain recordings ^31,32,80^, we addressed these unanswered questions. We found that the retrosplenial cortex was indeed tuned to changes in heading angle and, to a lesser extent, gaze activity (see figures 2C and 4C). Moreover, when distinguishing head direction from head rotation, inferior portions of the retrosplenial cortex appeared to be specifically tuned to head direction within an environment (see figure 6B). These results further expand our understanding of retrosplenial function in active navigation.

Combining intracranial recordings in patients with real-world navigational tasks is an even greater challenge than in healthy participants wearing scalp EEG, and very few studies have thus far successfully used the combination to investigate spatial tuning of human brain activity ^80^. At the same time, this rare combination opens up exciting opportunities to investigate human electrophysiology that would otherwise remain concealed ^28^ In the present study, the iEEG data allowed us to zoom in on the subregions of the medial temporal lobe, which is hardly possible with scalp EEG. Thereby, we were able to test and confirm predictions derived from animal models ^41^ and show that directional tuning is strong outside but not in the human hippocampus. On a more general level, we think of this approach as an important step towards more naturalistic settings in neuroscience, in particular for the fields of navigation and memory ^28^.

### On the influence of electromyographic (EMG) artifacts

Given the sensitivity of EEG recordings to EMG artifacts, one is right to question whether the results we report here are driven by EMG activity. We would suggest that this is not the case for five reasons:

1. Using a combination of ICA, PCA and LMEs on the scalp EEG data, we modelled EMG activity as a statistically independent effect from our main regressors of interest, minimising EMG influence on the central effect.
2. EMG effects could not explain why the scalp EEG activity best predicted head orientation when considering EEG activity ∼80ms before head rotation; if the central effect were attributable to EMG, the link between EEG and head orientation should be instantaneous. In the same vein, EMG effects would not be able to explain why scalp EEG sensitivity to head orientation persisted up to 500ms after the head rotation was completed.
3. EMG effects could not explain why the strongest scalp-level effects were observed at Pz, an electrode in the EEG cap that is one of the furthest away from any source of EMG activity (which arises most commonly around the mastoids and other electrodes at the edge of the cap).
4. In the source-based analysis of the scalp EEG data, we used LCMV beamformers which will project any EMG activity to outside the brain, meaning any sources located within the brain are unlikely to reflect an EMG effect.
5. In the intracranial analysis, we used bipolar re-referencing which will subtract out any common signals between two neighbouring electrodes (including EMG), again meaning that any observed effect is unlikely to reflect an EMG effect.

In any EEG study, even in those with fixed head position, it is difficult, if not impossible, to claim with absolute certainty that there are, emphatically, no EMG effects in the final results and we don’t set out to make such a claim here. We do however suggest that, because of the reasons outlined above, any muscular effect would be minimal and, as such, the effects observed here are more easily ascribed to a neural process rather than muscular one.

### Summary

In conclusion, we find electrophysiological signatures of veridical head direction in population-level activity of humans that shows a strong correspondence with head direction cell activity reported in rodents. By combining physical head rotations with a series of forward encoding models, we are able to provide a unique insight into the role of vestibular movements in human spatial navigation, overcoming a key limitation that has plagued such study to date ^28–30^, and revealing a detailed taxonomy of human head direction-related signals.

## Methods

All research complies with all relevant ethical regulations. The study was approved by the local ethics committee at Ludwig-Maximilians-Universität München.

### Participants

In Experiment 1, 39 healthy participants were recruited (mean age = 24.8 years, 61.5% female). They received course credit or financial reimbursement in return for their participation. All participants gave written informed consent after being informed about the details of the study. The study was approved by the local ethics committee. Due to recording difficulties stemming from the EEG and motion tracking systems, data from some tasks of some participants was lost. Table 1 contains a breakdown of which participants were excluded from which tasks. For the intracranial recordings, 10 patients undergoing treatment for medication-resistant epilepsy were recruited. The patients, who volunteered to participate in the study, had depth electrodes implanted for diagnostic reasons. The study was approved by the ethics committee of the Medical Faculty of the Ludwig–Maximilian Universität. All patients gave written informed consent after being informed about the details of the study. Difficulties with EEG/motion tracking also resulted in the exclusion of participants (see Table 1). In experiment 2, 24 healthy participants were recruited (mean age = 24.7, 54.2% female). They too received course credit or financial reimbursement in return for their participation and gave written informed consent after being informed about the details of the study.

**Table 1.**
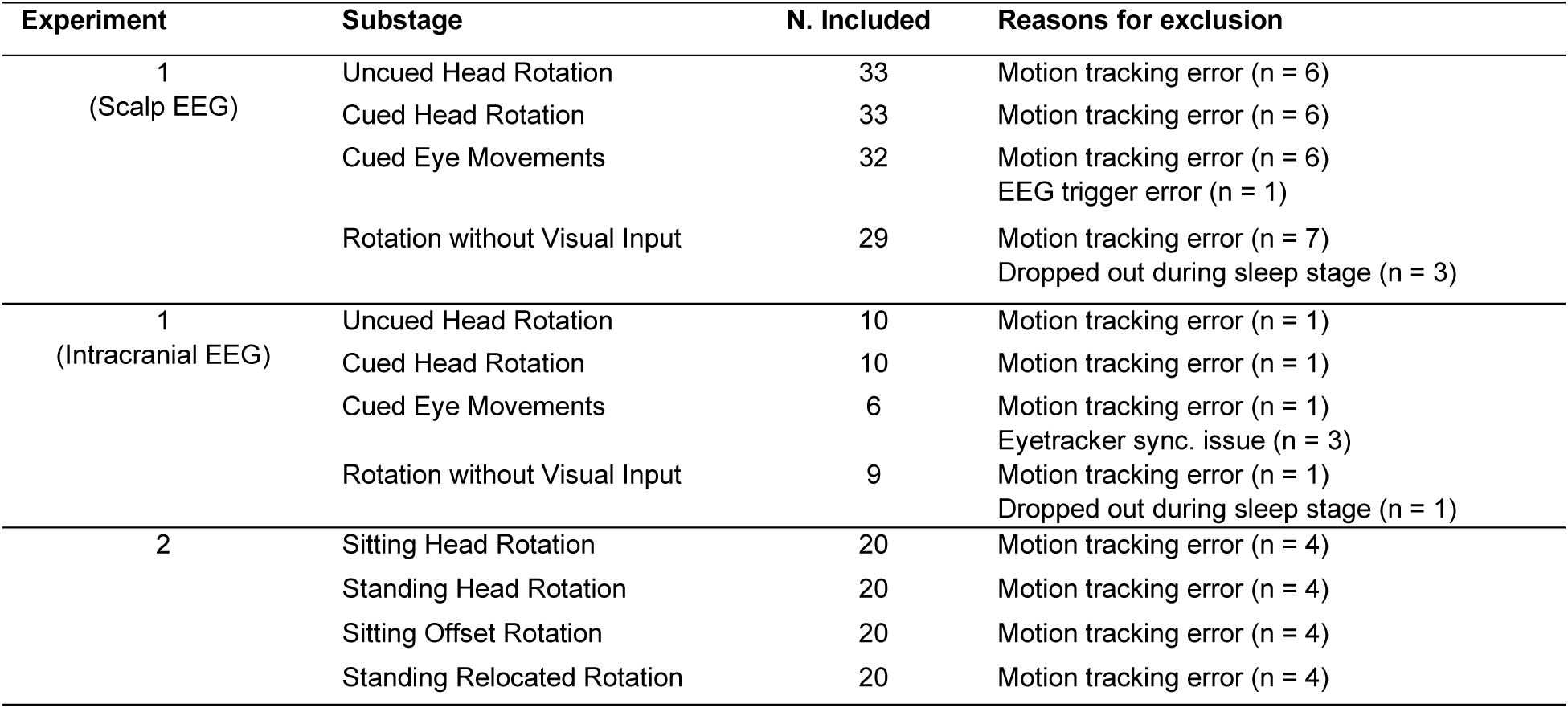
A list of the number of participants for a given task and the reasons for those who were excluded.

### Experimental design: Experiment 1

Participants arrived at the lab at approximately 7pm in the evening, where they were first fitted with an EEG cap, and then tasked with a series of experiments. At around 11pm, they slept in the lab, and woke at approximately 7am for a second series of experiments. These experiments can be broadly split into two categories: navigation and memory, both of which happened in the evening session and in the morning session. As this paper only relates to the navigation experiments, only the details of these tasks will be described.

Each navigation experiment followed the same typical pattern, delivered using Psychtoolbox (http://psychtoolbox.org/). Participants began by fixating on the centre screen. In all experiments, vertical screen positions match participants’ line of sight. A stimulus was presented that they had to respond to by turning to one of four other screens (positioned at -60°, -30°, +30° and 60° relative to the centre screen, see figure 1a). Each experiment consisted of 160 trials, split across 4 blocks (i.e., 40 trials per block). Below, each variation of this experiment is outlined in the order they occurred:

1. *Uncued Head Rotation:* At stimulus onset, the fixation cross disappeared and appeared on one of the four flanking screens. Participants were instructed to turn their head to face the screen which the fixation cross appeared on and fixate upon the cross. No auditory cue was played.
2. *Cued Head Rotation:* At stimulus onset and as the fixation moved to one of the four flanking screens, one of four animal sounds were played. Each animal sound was associated with a single flanking screen. Participants were instructed to turn their head to face the screen which the fixation cross appeared on and fixate upon the cross.
3. *Cued Eye Movement:* Programmatically speaking, this experiment matched the *Cued Head Rotation* exactly. However, participants received different instructions. Specifically, they were instructed to saccade to whichever flanking screen the fixation cross appeared on and fixate upon the cross while keeping their head facing towards the centre screen.
4. *Rotation without Visual Input:* As with the *Cued Eye Movements* experiment, programmatically speaking, this experiment matched the *Cued Head Rotation* exactly. In this case however, participants wore cardboard virtual reality goggles where a fixation cross was presented directly in front of them. When the auditory cue was played, they were instructed to turn their head to face the screen which they believed was associated with the auditory cue, all the while fixating on the cross presented via the virtual reality goggles. Note that while the previous three tasks were conducted in the evening, prior to sleep, this task was conducted in the morning. This was done to avoid tiring the participant out before they took part in the evening memory task.

### Experimental design: Experiment 2

This experiment focused solely on navigation with all tasks being completed in a single afternoon session. As in Experiment 1, each navigation task followed the same pattern. Participants fixated on the centre screen, before an auditory stimulus was presented that they had to respond to the associated screen (indicated by an animal image attached to the screen). In addition to the five screens present in Experiment 1 (see figure 1), a further two screens were added at 90° and 120° relative to the centre screen of Experiment 1. These two screens were associated with two additional animal sounds. These additional screens were only used in the third task. Each experiment consisted of 160 trials, split across 4 blocks (i.e., 40 trials per block). In all experiments, vertical screen positions match participants’ line of sight. Below, each variation of this experiment is outlined:

1. *Cued Head Rotation:* This was a direct replication of the *Cued Head Rotation* task used in Experiment 1.
2. *Standing Head Rotation:* This task matched the *Cued Head Rotation* task with the single exception that participants stood instead of sat. Participants were instructed to rotate their head while keeping their body facing forwards.
3. *Cued Head Rotation at 60° offset:* This task matched the *Cued Head Rotation* task with the single exception that participants sat facing the screen positioned at 60° from the centre screen and were then tasked with rotating their head relative to this position (that is, they rotated to face the screens at 0°, 30°, 90° and 120°).
4. *Standing Head Rotation after relocation:* This task matched the *Standing Head Rotation* task. Here however, participants stood ∼1 metre closer to the centre screen.

Note that, in this experiment, the order of the tasks was randomised such that tasks 2 and 4 could occur in either order. Task 1, however, always came first to help participants acclimatise to the task and match the position of this task in Experiment 1. Task 3 was always the third task as it gave the participants a chance to recover from standing.

### Scalp EEG acquisition and pre-processing

The EEG was recorded using an EEGo EEG system (ANT Neuro Enschede, Netherlands) with 65 Ag/AgCl electrodes arranged in a 10/10 system layout (including left and right mastoids; using CPz as the reference and AFz as the ground). All electrode impedances were less than 20 kΩ prior to commencing the experiment. The sampling rate was set at 1,000Hz.

All EEG analyses were conducted using the Fieldtrip toolbox 81 https://www.fieldtriptoolbox.org/) in conjunction with custom code. In preparation for the central analyses, the continuous EEG data was first high-pass (0.5Hz; Butterworth infinite impulse response [IIR]), low-pass (165Hz; Butterworth IIR) and band-stop filtered (49-51Hz; 99-101Hz; 149-151Hz; Butterworth IIR). Second, independent components analysis was conducted. In Experiment 1, we focused on only removing independent components that resembled eye blinks or saccades to preserve as much of the raw signal as possible. In Experiment 2, we took a more stringent approach and, in addition to removing eye blinks and saccades, removed probable electromuscular components. The strategy used did not impact the analysis outcome, as evidenced by the replication of the main effect across experiments (N.B., the more stringent approach used in Experiment 2 however may explain why the electromuscular regressor explained less variance in the linear mixed-effects model than it did in the first experiment). Third, the data was epoched around the onset of the auditory cue; these epochs began 1,500ms before the onset of the cue, and ended 4,500ms after the onset of the cue. In the case of the *Uncued Head Rotation* condition (in which no auditory cue was presented), the data was epoched around when the auditory cue would have, theoretically, occurred. Fourth, the data was visually inspected for artefactual trials and channels. Trials were removed when artifacts were temporally-limited and spread across many channels, whereas channels were removed when artifacts were present within a single channel for many trials (for examples of artifactual trials, see supplementary figure 13). Fifth, the artefactual channels that had been removed from the data in the previous step were interpolated using neighbouring channels. Sixth, the data was re-referenced using the average of all channels.

### iEEG acquisition and pre-processing

Recordings were performed at the Epilepsy Center, Department of Neurology, University of Munich, Germany. Intracranial EEG was recorded from Spencer depth electrodes (Ad-Tech Medical Instrument, Racine, Wisconsin, United States). Electrodes carried up to 15 contacts with, distances of 5 or 10mm between contacts. Data were recorded using XLTEK Neuroworks software (Natus Medical, San Carlos, California, US) and an XLTEK EMU128FS amplifier, with voltages referenced to a parietal electrode site. The sampling rate was set at 1,000Hz.

For signal preprocessing, the continuous iEEG data was first high-pass (0.5Hz; Butterworth IIR), low-pass (165Hz; Butterworth IIR) and band-stop filtered (49-51Hz; 99-101Hz; 149-151Hz; Butterworth IIR), identical to the scalp EEG preprocessing. Second, the data was epoched in the same manner as the scalp EEG data. Third, the data was then visually inspected for artefactual trials, in particular for interictal epileptic activity, and channels which exhibited such activity were marked and then removed from the data. Fourth, the data was re-referenced using a bipolar referencing scheme.

We estimated the locations of the intracranial contacts using the Lead-DBS software 82. First, we co-registered the post-operative CT scan to pre-operative T1-weighted image using a two-stage linear registration (rigid followed by affine) as implemented in Advanced Normalisation Tools 83. Second, we manually recorded the native space co-ordinates of all electrodes for the given patient with a specific focus on identifying and labelling those electrodes which sat in subregions of medial temporal lobe. Third, we spatially normalised these scans to MNI space based on the pre-operative T1-weighted image using the Unified Segmentation Approach as implemented in SPM12 84. Fourth, we manually recorded the MNI space co-ordinates of all electrodes for the given patient. Fifth, we re-labelled the contacts based on the lobe they sat in (as determined by the contact MNI co-ordinate and the lobe atlas available in *WFU_PickAtlas*; https://www.nitrc.org/projects/wfu_pickatlas/). Any contacts deemed to be within the medial temporal lobe were given the label of the visually identified subregion. See Supplementary Table 3 for a summary of electrodes per region-of-interest and per patient.

### Eye-tracking acquisition and pre-processing

Eye-tracking data was acquired using a Tobii Pro Spectrum system. This system was calibrated immediately prior to the beginning of the experiment. The sampling rate was set as 600Hz.

Offline, the co-ordinate space data was smoothed using a sliding window average of 50ms. The data was then converted to visual angles, epoched to auditory cue onset (using the same parameters as the EEG data) and visually inspected for artifacts. Any trial which contained physiologically-implausible changes in eye position were excluded (for an example trial, see supplementary figure 14). Moreover, any trial which was missing more than 25% of sample points were excluded. Any missing data points that remained were linearly interpolated based on data from the preceding and proceeding sample points. Note that only data from the three screens (center screen +/- 30°) were used as the eye-tracker could not reliably track eye positions on the outside screens. Lastly, the data was upsampled to match the EEG recording sample rate. Note that this upsampling procedure does not impact any lag-based analyses as the interpolated samples are drawn between two recorded samples spaced ∼2ms apart while the lag model slides the data in increments of 10ms, meaning the interpolated window is too short to spread across the sliding windows.

### Head motion-tracking acquisition and pre-processing

Head rotations were recorded using a Polhemus Liberty system. A sensor was attached the EEG cap and sat between the electrodes AFz and FPz. The reference sensor was placed ∼75cm behind and to the right of the participant. Similar positioning was used for the patients in Experiment 1, though some variability was encountered due to the position of the patient’s bandages. Polhemus’s proprietary software was used to record the change in position of the first sensor relative to the second. The sampling rate was set as 240Hz.

Offline, the co-ordinate space data was epoched around the onset of the auditory cue (as done for the EEG and eyetracking data). No smoothing or filtering was applied to the head motion data. The heading angle of every sample was operationalised as the yaw measurement (i.e., the rotation around the z-axis). The resulting head direction signal was then visually inspected for artefacts. Any trial which contained physiologically-implausible changes in head angle was excluded. Lastly, the data was upsampled to match the EEG recording sample rate. Note that this upsampling procedure does not impact any lag-based analyses as the interpolated samples are drawn between two recorded samples spaced ∼4ms apart while the lag model slides the data in increments of 10ms, meaning the interpolated window is too short to spread across the sliding windows.

### Forward encoding model: Overview

Here, we provide a brief overview of the forward encoding model (FEM) approach and, below, we report the specifics of each step. For a visualisation of the approach, see figure 7. The FEM we used took inspiration from that used by 27. First, we decomposed the real-time head angle measurement into basis sets of circular–Gaussian von-Mises distributions (from here on termed “HD kernels”). Second, taking these HD kernels and the recorded EEG voltage, we estimated an electrode’s directional tuning (that is, the relative weighting of each HD kernel for a given EEG electrode) using ridge regression. Finally, we applied these weights to held-out datasets from the same participant to test the generalisability of the model.

### Forward encoding model: Generating the model

We built a FEM for every participant, task, kernel width, and EEG electrode individually. Head angle was modelled using a basis set of circular–Gaussian von-Mises distributions (as implemented in the Circular Statistics Toolbox for Matlab; https://github.com/circstat/circstatmatlab), with total kernel space covering 180° (i.e., ±90° from the centre screen) with an angular resolution of 1°. To test the tuning width of each electrode, we iterated our analysis across six models, which each used a bespoke kernel width (6°, 10°, 15°, 20°, 30°, 45°, and 60°; matching 27). The distance between each kernel was tied to the kernel width to control for directional sensitivity across models (i.e., models with a large kernel width used fewer kernels). Note that as the kernels overlap, heading angles that sit between the peaks of two kernels can be modelled as the sum of the two kernels. We then estimated the magnitude of kernel activities for every time point based on the real-time head angle. By doing this for all kernels and time points, we obtained a 2D matrix (termed ‘X’) that described head angle as a function of time and directional tuning.

### Forward encoding model: Training the model

The motion signal and EEG voltage were split into four partitions; each partition contained the same number of trials for each head rotation. The training dataset was then formed from three of these partitions, and the testing dataset was the remaining partition. This was done in a cross-validated manner such that, through iteration, each partition acted as the testing dataset for the other partitions. Note that simulations indicate that the pre-selection of target head angles and the skewed distribution this produces in sampling does not bias the model (see supplementary figure 15).

To train the FEM, we used the matrix derived in the section above (X) in conjunction with the EEG training data at a given electrode (y) to predict the HD kernel weightings (β) through ridge (i.e., L2-regularised) regression. Ridge regression helps penalise excessively high coefficients that may come about through the fact that the HD kernels for a given timepoint are not independent. However, the regularisation parameter (λ) for this approach first needs to be estimated. To do this, we used cross-validation; we took two of the three training partitions, estimated the HD kernel weightings (β) for ten different values of λ (log-spaced between 1 and 10,000,000), and then assessed how well these values generalised to the remaining training partition. To assess generalisability, we calculated the Pearson correlation coefficient between the time course of the EEG voltage and the time course predicted by the β-weighted HD kernels for each value of λ. The value of λ that led to the highest Pearson correlation coefficient (averaged across the cross-validations) was then used to estimate the final model weights. The final model weights were derived using ridge regression (this time across all three partitions).

The stability of the weights and λ are visualised separately for each experiment (see supplementary figures 16-22).

### Forward encoding model: Testing the model

To test the validity of the FEM, we applied the model weights to head angle data held in the independent testing dataset (one out of four partitions). This returned a prediction of the EEG time-series, which was then correlated with the observed EEG voltage time-series using Pearson’s correlation co-efficient.

To aid comparison between the models using different kernel widths, we used a bootstrapping procedure where we circularly-shifted the predicted EEG voltage time-series by a random integer and then recomputed the correlation coefficient. By repeating this procedure 100 times, we generated a distribution of correlation coefficients between the observed and predicted time-series that we would expect by chance. We can then normalise the “true” correlation coefficient by subtracting the mean of the chance distribution and then dividing by the standard deviation of this distribution.

### Forward encoding model: Modelling muscular activity

Muscular activity was modelled as a covariate in the linear mixed-effects (LME) model used for inferential analysis, and hence regressed out of the variables of interest. The full model is described below (see “*EEG encoding model: Group-level analysis for Experiment 1”*). Here, we just describe the component of the LME model used to regress out muscular activity.

To this end, we used principal components analysis (PCA) to break the EEG data into its principal components, and then ordered these components based on the relative likelihood that a given component contained muscular activity, informed by the components’ topography. This likelihood measure was computed by taking the mean of the size of the weights for electrodes on the outer rim of the cap and dividing this value by the mean of the size of the weights for all the remaining electrodes. The larger the resulting value, the more ‘muscular’ a component was judged to be, following the rationale that electrodes that sit at the edge of the cap are closest to the neck muscles involved in the head rotation. Based on this ordering procedure, we created five groups of components: the 20% least muscular components, the 40% least muscular components, and so on. For each of these groups, the EEG data was reconstructed in electrode-space using the selected components. The FEM was generated, trained, and tested for each of these datasets separately.

The performance of these FEMs can be thought of as being a linear sum of EEG signals that reflect muscular activity and EEG signals that do not reflect muscular activity (there can be no third option – the signal is either muscular or not). Based on the PCA approach used above, we can also conclude that the EEG signals that reflect muscular activity get progressively smaller as we remove more of the muscular components, while the EEG signals that do not reflect muscular activity remain constant. This can be expressed as:

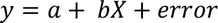

Where *y* represents FEM performance, *a* represents the constant (i.e., EEG signal that doesn’t change as a function of the number of muscular components removed), *b* represents an unknown weighting, and *X* reflects the percentage of muscular components removed. This equation will form part of the central LME model and allow for muscular contributions to FEM performance to be regressed out of the central analyses.

### Forward encoding model: Modelling eye-position activity

In addition to creating a FEM which used head angle as a predictor of EEG activity, we also created a FEM which used eye position as a predictor of EEG activity. The eye position FEM was computed in the same manner as the head angle FEM with two exceptions: (1) visual angle was used as the predictor in place of head angle, and (2) only data from the screens as +/- 30° were used as the eye-tracker could not track eye positions on the outside screens. The eye position FEM was built on the data from the *Cued Eye Movements* task. As no head rotation occurred in this task, whenever we refer to FEM performance for this task, we refer to the eye position model.

### Forward encoding model: Group-level analysis of Experiment 1

Through completion of the analysis steps described above, we obtained a z-score describing FEM performance relative to chance for every participant, task, electrode, kernel width and principal component subcategory. To assess whether these z-scores differed from chance across participants (i.e., z > 0), we used linear mixed-effect (LME) models. We computed an LME for every kernel-electrode pair. Each LME took data from all participants, all tasks and all principal components subcategories. The outcome variable of each LME was FEM performance. For predictors, random intercepts were created for each participant, as well as four fixed effects that represented latent factors. The fixed effects were as follows:

1. Head angle: A binary variable which denoted whether head rotation had occurred in the given task.
2. Auditory input: A binary variable which denoted whether an auditory cue had been played in the given task.
3. Visual input / eye movements: A binary variable which denoted whether visual input which, in these experiments, occurred whenever participants were free to move their eyes.
4. Muscular activity: A variable ranging from 0.2 to 1, which denoted the percentage of principal components included in the sample prior to the fitting of the encoding model.

We felt it was essential to simultaneously model and, consequently, factor out the confounds of auditory input, visual input/eye movements and muscular activity as these confounds co-occurred with the head rotations and it would be entirely plausible to suggest that the ability of the FEM to predict head orientation was driven not by head direction per se, but by neural responses to auditory cues telling participants where to turn to, changes in visual input that accompany the change in head direction, or muscle activity required to rotate the head. See Supplementary Table 4 for a mapping of experiment conditions onto these latent factors.

The LME returned beta weights for each regressor. To assess statistical significance, the resulting weights were compared to a surrogate distribution (200 permutations) of weights generated by randomly permuting the sign of the outcome variable for each sample and then refitting the model. The p-value was computed by calculating the proportion of surrogate weights which were larger in value than the ‘true’ model weights (this is equivalent to a one-tailed test). This process was repeated for every kernel-electrode pair. False discovery rate (FDR) correction was used to address the multiple comparison issue incurred by conducting analysis on each resulting beta weight, kernel and electrode separately.

The LME used for the patients was near-identical to that used for the healthy participants. The key difference was that, rather than using separate models for each electrode, separate models were used for defined regions-of-interest. This approach was necessitated by the fact that implantation montages are bespoke for each patient, leading to a variable number of contacts in each region across patients. Regions-of-interest were defined as the frontal lobe, occipital lobe, parietal lobe, and temporal lobe (as defined by the lobe options in WFU_PickAtlas; https://www.nitrc.org/projects/wfu_pickatlas/), and the parahippocampus, hippocampus and amygdala (as defined by visual inspection of the patient scans. As before, participants were included as random effects in the LME, avoiding issues in generalisability that occur when using fixed-effect models.

### Forward encoding model: Group-level analysis of Experiment 2

The concept behind the group-level analysis for Experiment 2 matched that of Experiment 1, but given the different tasks, the latent factors inevitably changed. Here, the fixed effects were as follows:

1. Head angle: A binary variable which denoted whether head rotation had occurred in the given task.
2. Position on the z-axis: A binary variable which denoted whether participants were sitting or standing during the given task.
3. Relocation on the x-axis: A binary variable which denoted whether participants were sitting/standing in the standard position during the given task, or whether they stepped one metre closer to the screens.
4. Muscular activity: A variable ranging from 0.2 to 1, which denoted the percentage of principal components included in the data prior to the fitting of the encoding model.

See Supplementary Table 5 for a mapping of experiment conditions onto latent factors.

### Lag-based encoding model

To assess the temporal dynamics of the EEG signal tuned to head angle relative to the timing of physical head rotation, we built a lag-based FEM. This FEM operates in the same manner as described above, except that (1) the time series of the EEG is shifted by a given number of samples prior to fitting, and (2) to reduce computational load, only kernels of 20° were used. We restricted the analysis to the 20° kernel FEM as this FEM best fit the EEG data. Note that as we are interested in which lag the FEM operates at best, rather than simply whether the FEM operates above chance, circularity is a non-issue as all lags benefit from the same kernel selection process. The temporal shift of the EEG is referred to as the lag. We fitted the lag-based model for 40 lags, ranging from when the EEG time series preceded the motion tracking time series by 200ms to when the EEG time series followed the motion tracking time series by 200ms. All other aspects of the analysis remained the same. False discovery rate (FDR) correction was used to address the multiple comparison issue incurred by conducting analysis on each electrode and lag separately.

To identify whether the lag-based encoding model performed better when EEG activity led or lagged heading angle, the difference was computed between the mean of LME beta coefficients for “leading” samples and the mean of LME beta coefficients for “lagging” samples. This was done for the coefficients derived from the true, observed data, and for coefficients derived from the permuted “chance” data. The resulting difference measure for the true data was compared to the permuted distribution, and a both a z-value and a p-value was computed. The z-value used the mean and standard deviation of the permutation distribution, while the p-value counted the percentage of chance data points that had a larger difference score than the observed data.

### Inverted encoding model

As a second means of assessing the temporal dynamics of the head angle-related EEG signal, we build an inverted encoding model. This approach matched the main approach up until the section “*EEG encoding model: Testing the model*”. At this stage, the weights derived from 20° FEM were inverted (i.e., *inv*(*β* ∗ *β*’) ∗ *β*). The weights were then applied to the EEG scalp data iteratively for each sample point. This returned predicted head direction kernel values for every sample of every trial. Representational similarity analysis (RSA) was then used to identify whether the kernel magnitudes at a given sample could distinguish between the four monitor positions. To this end, a representational dissimilarity matrix (RDM) was computed by computing Pearson correlations between the predicted kernel magnitudes of every pair of trials. The RDM was then correlated with a model RDM which stated that the correlation between pairs with the same target head direction would equal one and all other pairs would equal zero. A surrogate distribution of chance correlation values was then computed by shuffling the labels of the trials, rebuilding the model RDM and then correlating the shuffled model RDM with the observed RDM. The correlation of the ‘true’ data was then normalised using the mean and standard deviation of the surrogate data. By repeating these steps for every time point, we obtain a time series that describes the extent to which the inverted model predicts head direction above and beyond what would be expected by chance. False discovery rate (FDR) correction was used to address the multiple comparison issue incurred by conducting analysis on each time point separately. The remainder of the analysis matches that of the main approach.

### Generalised encoding models

To disentangle the influences of head rotation and head direction signal, and location-specific and location-independent head direction signal, we used generalised encoding models. We will use the head rotation/head direction dissociation as the example here, but the same procedure was applied to the location-specific/location-independent dissociation.

For the generalised encoding models, we trained the encoding model on one condition (e.g., Cued Head Rotation) and applied the model weights to another condition (i.e., Cued Head Rotation at 60° offset). In the case of head direction versus head rotation, if the weights lead to the test data outperforming what would be expected by chance, it can be said that the model tracks activity related to head rotation as the FEM generalises to when participants are sitting at an offset. If the weights, however, lead to poorer performance than what is observed when conducting cross-validated training and testing within a condition, it can be said that the model tracks activity head direction. Notably, the two outcomes are not mutually exclusive: generalised model performance can be significantly greater than chance, while also being significantly poorer than within-condition performance. To test both ideas simultaneously, linear mixed models were used with fixed effects denoting head rotation, head direction and muscular activity (see Supplementary Table 6 for mapping of generalised encoding models onto factors). Random intercepts were included for each participant. Statistical analysis matched what was reported above.

The same principles were applied to the dissociation of location-independent and location-specific head direction signals. Supplementary Table 7 describes how the generalised encoding models were mapped onto the fixed effects of the linear mixed effect models.

## Code availability

Openly available at https://github.com/benjaminGriffiths/human-hd.

## Data availability

Data acquired from the healthy participants is available at https://data.ub.uni-muenchen.de/439/. Due to privacy laws, data acquired from the patient is not openly available, though (subject to privacy laws) can be provided by contacting the corresponding author.

## Acknowledgements

This work was supported by the European Research Council (https://erc.europa.eu/, Starting Grant 802681 awarded to TSt) and the Leverhulme Trust (https://www.leverhulme.ac.uk/, Early Career Fellowship ECF-2021-628 awarded to BJG). The funders had no role in study design, data collection and analysis, decision to publish or preparation of the manuscript. We thank all participants and in particular all patients who volunteered to participate in this study. We thank the staff and physicians at the Epilepsy Center, Department of Neurology, Ludwig-Maximilians-Universität, Munich, for assistance. We thank Aditya Chowdhury for valuable input.

## Author Contributions Statement

BJG: conceptualization, methodology, software, validation, formal analysis; investigation; data curation; writing – original draft; writing – review & editing; visualization; ThS: conceptualization, methodology, investigation, data curation, writing – review & editing; JS: investigation, data curation, writing – review & editing; CV: resources, writing – review & editing; EK: resources, writing – review & editing; SQ: resources, writing – review & editing; JR: resources, writing – review & editing; SN: resources, writing – review & editing; ToS: conceptualization, methodology, investigation; writing – original draft; writing – review & editing, supervision, project administration, funding acquisition.

## References

1. Buzsáki, G. & Moser, E. I. Memory, navigation and theta rhythm in the hippocampal-entorhinal system. Nat. Neurosci. 16, 130–138 (2013).

2. Baumann, O. & Mattingley, J. B. Extrahippocampal contributions to spatial navigation in humans: A review of the neuroimaging evidence. Hippocampus 1–18 (2021) doi:10.1002/hipo.23313.

3. Bellmund, J. L. S., Gärdenfors, P., Moser, E. I. & Doeller, C. F. Navigating cognition: Spatial codes for human thinking. Science 362, (2018).

4. Epstein, R. A., Patai, E. Z., Julian, J. B. & Spiers, H. J. The cognitive map in humans: Spatial navigation and beyond. Nat. Neurosci. 20, 1504–1513 (2017).

5. Geva-Sagiv, M., Las, L., Yovel, Y. & Ulanovsky, N. Spatial cognition in bats and rats: from sensory acquisition to multiscale maps and navigation. Nat. Rev. Neurosci. 16, 94–108 (2015).

6. Hulse, B. K. & Jayaraman, V. Mechanisms underlying the neural computation of head direction. Annu. Rev. Neurosci. 31–54 (2019) doi:10.1146/annurev-neuro-072116-031516.

7. Long, X. & Zhang, S. J. A novel somatosensory spatial navigation system outside the hippocampal formation. Cell Res. 1–15 (2021) doi:10.1038/s41422-020-00448-8.

8. Muller, R. U., Ranck, J. B. & Taube, J. S. Head direction cells: properties and functional significance. Curr. Opin. Neurobiol. 6, 196–206 (1996).

9. Ranck, J. B. Head direction cells in the deep layer of dorsal presubiculum in freely moving rats. in Soc. Neuroscience Abstr. vol. 10 599 (1984).

10. Sharp, P. E., Blair, H. T. & Cho, J. The anatomical and computational basis of the rat head-direction cell signal. Trends Neurosci. 24, 289–294 (2001).

11. Taube, J. S. The Head Direction Signal: Origins and Sensory-Motor Integration. Annu. Rev. Neurosci. 30, 181–207 (2007).

12. Taube, J. S. & Bassett, J. P. Persistent Neural Activity in Head Direction Cells. Cereb. Cortex 13, 1162–1172 (2003).

13. Taube, J. S., Muller, R. & Ranck, J. Head-direction cells recorded from the postsubiculum in freely moving rats. I. Description and quantitative analysis. J. Neurosci. 10, 420–435 (1990).

14. Taube, J. S. Head Direction Cells Recorded in the Anterior Thalamic Nuclei of Freely Moving Rats. J. Neurosci. 17 (1995).

15. Taube, J. S. & Muller, R. U. Comparisons of head direction cell activity in the postsubiculum and anterior thalamus of freely moving rats. Hippocampus 8, 87–108 (1998).

16. Goodridge, J. P., Dudchenko, P. A., Worboys, K. A., Golob, E. J. & Taube, J. S. Cue Control and Head Direction Cells. Behav. Neurosci. 112, 749–761 (1998).

17. Blair, H. T. & Sharp, P. Anticipatory head direction signals in anterior thalamus: evidence for a thalamocortical circuit that integrates angular head motion to compute head direction. J. Neurosci. 15, 6260–6270 (1995).

18. Zirkelbach, J., Stemmler, M. & Herz, A. V. M. Anticipatory Neural Activity Improves the Decoding Accuracy for Dynamic Head-Direction Signals. J. Neurosci. 39, 2847–2859 (2019).

19. Butler, W. N., Smith, K. S., van der Meer, M. A. A. & Taube, J. S. The head-direction signal plays a functional role as a neural compass during navigation. Curr. Biol. 10 (2017).

20. Calton, J. L. et al. Hippocampal Place Cell Instability after Lesions of the Head Direction Cell Network. J. Neurosci. 23, 9719–9731 (2003).

21. Winter, S. S., Clark, B. J. & Taube, J. S. Disruption of the head direction cell network impairs the parahippocampal grid cell signal. Science 347, 870–874 (2015).

22. Harland, B. et al. Lesions of the Head Direction Cell System Increase Hippocampal Place Field Repetition. Curr. Biol. 27, 2706–2712.e2 (2017).

23. Ajabi, Z., Keinath, A. T., Wei, X.-X. & Brandon, M. P. Population dynamics of head-direction neurons during drift and reorientation. Nature 615, 892–899 (2023).

24. Chaudhuri, R., Gerçek, B., Pandey, B., Peyrache, A. & Fiete, I. The intrinsic attractor manifold and population dynamics of a canonical cognitive circuit across waking and sleep. Nat. Neurosci. 22, 1512–1520 (2019).

25. Buzsáki, G., Anastassiou, C. A. & Koch, C. The origin of extracellular fields and currents-EEG, ECoG, LFP and spikes. Nat. Rev. Neurosci. 13, 407–420 (2012).

26. Kunz, L. et al. Mesoscopic Neural Representations in Spatial Navigation. Trends Cogn. Sci. 23, 615–630 (2019).

27. Nau, M., Navarro Schröder, T., Frey, M. & Doeller, C. F. Behavior-dependent directional tuning in the human visual-navigation network. Nat. Commun. 11, 1–13 (2020).

28. Stangl, M., Maoz, S. L. & Suthana, N. Mobile cognition: imaging the human brain in the ‘real world’. Nat. Rev. Neurosci. 24, 347–362 (2023).

29. Steel, A., Robertson, C. E. & Taube, J. S. Current Promises and Limitations of Combined Virtual Reality and Functional Magnetic Resonance Imaging Research in Humans: A Commentary on Huffman and Ekstrom (). J. Cogn. Neurosci. 33, 159–166 (2021).

30. Taube, J. S., Valerio, S. & Yoder, R. M. Is Navigation in Virtual Reality with fMRI Really Navigation? J. Cogn. Neurosci. 25, 1008–1019 (2013).

31. Do, T.-T. N., Lin, C.-T. & Gramann, K. Human brain dynamics in active spatial navigation. Sci. Rep. 11, 13036 (2021).

32. Gramann, K., Hohlefeld, F. U., Gehrke, L. & Klug, M. Human cortical dynamics during full-body heading changes. Sci. Rep. 11, 18186 (2021).

33. Griffiths, B. J., Mazaheri, A., Debener, S. & Hanslmayr, S. Brain oscillations track the formation of episodic memories in the real world. NeuroImage 143, 256–266 (2016).

34. Maoz, S. L. L. et al. Dynamic neural representations of memory and space during human ambulatory navigation. Nat. Commun. 14, 6643 (2023).

35. Piñeyro Salvidegoitia, M., et al. Out and about: Subsequent memory effect captured in a natural outdoor environment with smartphone EEG. Psychophysiology e13331–e13331 (2019) doi:10.1111/psyp.13331.

36. Schreiner, T. et al. Memory reactivation of real-world spatial orientation revealed by human electrophysiology. bioRxiv (2023) doi:10.1101/2023.01.27.525854.

37. Giocomo, L. M. et al. Topography of Head Direction Cells in Medial Entorhinal Cortex. Curr. Biol. 24, 252–262 (2014).

38. Wirth, S., Baraduc, P., Planté, A., Pinède, S. & Duhamel, J.-R. Gaze-informed, task-situated representation of space in primate hippocampus during virtual navigation. PLOS Biol. 15, e2001045 (2017).

39. Piza, D. B. et al. The hippocampus of the common marmoset is a GPS, but G is for gaze. bioRxiv (2023) doi:10.1101/2023.05.24.542209.

40. Kriegeskorte, N. Representational similarity analysis – connecting the branches of systems neuroscience. Front. Syst. Neurosci. 2, 1–28 (2008).

41. Cullen, K. E. & Taube, J. S. Our sense of direction: progress, controversies and challenges. Nat. Neurosci. 20, 1465–1473 (2017).

42. Bernardi, S. et al. The Geometry of Abstraction in the Hippocampus and Prefrontal Cortex. Cell 183, 954–967.e21 (2020).

43. Dudchenko, P. A., Wood, E. R. & Smith, A. Neurosci. Biobehav. Rev. 105, 24–33 (2019).

44. Patai, E. Z. & Spiers, H. J. The Versatile Wayfinder: Prefrontal Contributions to Spatial Navigation. Trends Cogn. Sci. 25, 520–533 (2021).

45. Jacob, P.-Y. et al. An independent, landmark-dominated head-direction signal in dysgranular retrosplenial cortex. Nat. Neurosci. 20, 173–175 (2017).

46. Lomi, E., Jeffery, K. J. & Mitchell, A. S. Convergence of direction, location and theta in the rat anteroventral thalamic nucleus. bioRxiv (2023) doi:10.1101/2023.01.11.523585.

47. Calton, J. L., Turner, C. S., Cyrenne, D.-L. M., Lee, B. R. & Taube, J. S. Landmark control and updating of self-movement cues are largely maintained in head direction cells after lesions of the posterior parietal cortex. Behav. Neurosci. 122, 827–840 (2008).

48. Medendorp, W. P., Beurze, S. M., Van Pelt, S. & Van Der Werf, J. Behavioral and cortical mechanisms for spatial coding and action planning. Cortex 44, 587–597 (2008).

49. Blair, H. T., Cho, J. & Sharp, P. E. Role of the Lateral Mammillary Nucleus in the Rat Head Direction Circuit: A Combined Single Unit Recording and Lesion Study. Neuron 21, 1387–1397 (1998).

50. Stackman, R. W. & Taube, J. S. Firing Properties of Rat Lateral Mammillary Single Units: Head Direction, Head Pitch, and Angular Head Velocity. J. Neurosci. 18, (1998).

51. Bassett, J. P. & Taube, J. S. Neural Correlates for Angular Head Velocity in the Rat Dorsal Tegmental Nucleus. J. Neurosci. 21, 5740–5751 (2001).

52. Shinder, M. E. & Taube, J. S. Self-motion improves head direction cell tuning. J. Neurophysiol. 111, 2479–2492 (2014).

53. Shine, J. P., Valdés-Herrera, J. P., Hegarty, M. & Wolbers, T. The Human Retrosplenial Cortex and Thalamus Code Head Direction in a Global Reference Frame. J. Neurosci. 36, 6371–6381 (2016).

54. Shinder, M. E. & Taube, J. S. Active and passive movement are encoded equally by head direction cells in the anterodorsal thalamus. J. Neurophysiol. 106, 788–800 (2011).

55. Laurens, J., Kim, B., Dickman, J. D. & Angelaki, D. E. Gravity orientation tuning in macaque anterior thalamus. Nat. Neurosci. 19, 1566–1568 (2016).

56. Robertson, R. G., Rolls, E. T., Georges-Francois, P. & Panzeri, S. Head direction cells in the primate pre-subiculum. Hippocampus 9, 206–219 (1999).

57. Georges-Francois, P. Spatial View Cells in the Primate Hippocampus: Allocentric View not Head Direction or Eye Position or Place. Cereb. Cortex 9, 197–212 (1999).

58. Julian, J. B., Keinath, A. T., Frazzetta, G. & Epstein, R. A. Human entorhinal cortex represents visual space using a boundary-anchored grid. Nat. Neurosci. 21, 191–194 (2018).

59. Killian, N. J., Jutras, M. J. & Buffalo, E. A. A map of visual space in the primate entorhinal cortex. Nature 491, 761–764 (2012).

60. Meister, M. L. R. & Buffalo, E. A. Neurons in Primate Entorhinal Cortex Represent Gaze Position in Multiple Spatial Reference Frames. J. Neurosci. 38, 2430–2441 (2018).

61. Nau, M., Navarro Schröder, T., Bellmund, J. L. S. & Doeller, C. F. Hexadirectional coding of visual space in human entorhinal cortex. Nat. Neurosci. 21, 188–190 (2018).

62. Rolls, E. T. Spatial view cells and the representation of place in the primate hippocampus. Hippocampus 9, 467–480 (1999).

63. Rolls, E. T. & O’Mara, S. M. View-responsive neurons in the primate hippocampal complex. Hippocampus 5, 409–424 (1995).

64. Staudigl, T. et al. Hexadirectional Modulation of High-Frequency Electrophysiological Activity in the Human Anterior Medial Temporal Lobe Maps Visual Space. Curr. Biol. 28, 3325–3329.e4 (2018).

65. Killian, N. J., Potter, S. M. & Buffalo, E. A. Saccade direction encoding in the primate entorhinal cortex during visual exploration. Proc. Natl. Acad. Sci. 112, 15743–15748 (2015).

66. Cowie, R. J. & Robinson, D. L. Subcortical contributions to head movements in macaques. I. Contrasting effects of electrical stimulation of a medial pontomedullary region and the superior colliculus. J. Neurophysiol. 72, (1994).

67. Elsley, J. K., Nagy, B., Cushing, S. L. & Corneil, B. D. Widespread Presaccadic Recruitment of Neck Muscles by Stimulation of the Primate Frontal Eye Fields. J. Neurophysiol. 98, 1333–1354 (2007).

68. Goonetilleke, S. C., Gribble, P. L., Mirsattari, S. M., Doherty, T. J. & Corneil, B. D. Neck muscle responses evoked by transcranial magnetic stimulation of the human frontal eye fields: TMS of the FEF evokes neck muscle activity. Eur. J. Neurosci. 33, 2155–2167 (2011).

69. Gu, C. & Corneil, B. D. Transcranial Magnetic Stimulation of the Prefrontal Cortex in Awake Nonhuman Primates Evokes a Polysynaptic Neck Muscle Response That Reflects Oculomotor Activity at the Time of Stimulation. J. Neurosci. 34, 14803–14815 (2014).

70. Sparks, D. L., Freedman, E. G., Chen, L. L. & Gandhi, N. J. Cortical and subcortical contributions to coordinated eye and head movements. Vision Res. 41, 3295–3305 (2001).

71. Baumann, O. & Mattingley, J. B. Medial Parietal Cortex Encodes Perceived Heading Direction in Humans. J. Neurosci. 30, 12897–12901 (2010).

72. Jacobs, J., Kahana, M. J., Ekstrom, A. D., Mollison, M. V. & Fried, I. A sense of direction in human entorhinal cortex. Proc. Natl. Acad. Sci. 107, 6487–6492 (2010).

73. Kim, M. & Maguire, E. A. Encoding of 3D head direction information in the human brain. Hippocampus 29, 619–629 (2019).

74. Kunz, L. et al. A neural code for egocentric spatial maps in the human medial temporal lobe. Neuron 109, 2781–2796.e10 (2021).

75. Vass, L. K. & Epstein, R. A. Abstract Representations of Location and Facing Direction in the Human Brain. J. Neurosci. 33, 6133–6142 (2013).

76. Stackman, R. W. & Taube, J. S. Firing Properties of Head Direction Cells in the Rat Anterior Thalamic Nucleus: Dependence on Vestibular Input. J. Neurosci. 17, 4349–4358 (1997).

77. Yoder, R. M. et al. Both visual and idiothetic cues contribute to head direction cell stability during navigation along complex routes. J. Neurophysiol. 105, 2989–3001 (2011).

78. Gramann, K., Ferris, D. P., Gwin, J. & Makeig, S. Imaging natural cognition in action. Int. J. Psychophysiol. 91, 22–29 (2014).

79. Lu, Z., Julian, J., B.,. & Epstein, R. A. Coding of head direction in the human visual system during dynamic navigation. J. Vis. 22, 4277–4277 (2022).

80. Stangl, M. et al. Boundary-anchored neural mechanisms of location-encoding for self and others. Nature 589, 420–425 (2021).

81. Oostenveld, R., Fries, P., Maris, E. & Schoffelen, J.-M. FieldTrip: Open source software for advanced analysis of MEG, EEG, and invasive electrophysiological data. Comput. Intell. Neurosci. 2011, 1–9 (2011).

82. Horn, A. & Kühn, A. A. Lead-DBS: A toolbox for deep brain stimulation electrode localizations and visualizations. NeuroImage 107, 127–135 (2015).

83. Avants, B. B., Epstein, C. L., Grossman, M. & Gee, J. C. Symmetric diffeomorphic image registration with cross-correlation: Evaluating automated labeling of elderly and neurodegenerative brain. Med. Image Anal. 12, 26–41 (2008).

84. Ashburner, J. & Friston, K. J. Unified segmentation. NeuroImage 26, 839–851 (2005).

