## Supplementary Materials for "Electrophysiological signatures of veridical head direction in humans"

for

**Supplementary Table 1.** Fixed effects for the basic linear mixed-effects model used in Experiment 1.

| Regressor | Tuning Width | Peak Electrode | z | $p_{FDR}$ |
|---|---|---|---|---|
| Head Direction | 6 | C3 | 6.457 | < 0.001 |
| | 10 | P1 | 6.272 | < 0.001 |
| | 15 | C5 | 5.753 | < 0.001 |
| | 20 | C5 | 6.711 | < 0.001 |
| | 30 | C5 | 5.951 | < 0.001 |
| | 45 | C5 | 6.488 | < 0.001 |
| | 60 | C5 | 6.629 | < 0.001 |
| Auditory Cue | 6 | CP3 | 0.700 | 0.414 |
| | 10 | CP3 | 0.632 | 0.471 |
| | 15 | CP3 | 0.855 | 0.377 |
| | 20 | CP3 | 0.925 | 0.361 |
| | 30 | CP3 | 1.402 | 0.226 |
| | 45 | CP3 | 1.538 | 0.178 |
| | 60 | Fpz | 1.274 | 0.268 |
| Visual Input / Eye-movements | 6 | Pz | 1.127 | 0.310 |
| | 10 | P6 | 0.624 | 0.488 |
| | 15 | Pz | 0.244 | 0.619 |
| | 20 | P6 | 1.002 | 0.361 |
| | 30 | C5 | 1.248 | 0.277 |
| | 45 | P6 | 1.273 | 0.217 |
| | 60 | C5 | 1.536 | 0.156 |
| Electromuscular activity | 6 | AF8 | 3.501 | < 0.001 |
| | 10 | AF8 | 3.957 | < 0.001 |
| | 15 | AF8 | 4.097 | < 0.001 |
| | 20 | AF8 | 3.773 | < 0.001 |
| | 30 | F8 | 4.035 | < 0.001 |
| | 45 | F7 | 4.060 | < 0.001 |
| | 60 | AF8 | 3.818 | < 0.001 |

**Supplementary Table 2.** Fixed effects for the basic linear mixed-effects model used in Experiment 2.

| Regressor | Tuning Width | Peak Electrode | z | $p_{FDR}$ |
|---|---|---|---|---|
| Head Direction | 6 | POz | 6.823 | < 0.001 |
| | 10 | Cz | 6.552 | < 0.001 |
| | 15 | Cz | 6.240 | < 0.001 |
| | 20 | Cz | 7.063 | < 0.001 |
| | 30 | FC2 | 6.277 | < 0.001 |
| | 45 | Cz | 6.746 | < 0.001 |
| | 60 | Cz | 7.548 | < 0.001 |
| Relocation x-axis | 6 | TP7 | 0.074 | 0.470 |
| | 10 | P4 | 0.568 | 0.325 |
| | 15 | P4 | 0.610 | 0.235 |
| | 20 | P4 | 0.865 | 0.225 |
| | 30 | TP7 | 0.747 | 0.250 |
| | 45 | T8 | 0.526 | 0.285 |
| | 60 | TP7 | 0.931 | 0.195 |
| Relocation z-axis | 6 | FCz | 1.788 | 0.065 |
| | 10 | FCz | 1.795 | 0.045 |
| | 15 | FCz | 1.914 | 0.030 |
| | 20 | CP5 | 1.405 | 0.090 |
| | 30 | F2 | 1.706 | 0.060 |
| | 45 | F2 | 1.688 | 0.050 |
| | 60 | AFz | 1.215 | 0.130 |
| Electromuscular activity | 6 | AF8 | 3.559 | < 0.001 |
| | 10 | FC5 | 3.342 | < 0.001 |
| | 15 | F5 | 3.516 | < 0.001 |
| | 20 | FC5 | 3.802 | < 0.001 |
| | 30 | F5 | 4.667 | < 0.001 |
| | 45 | F5 | 4.063 | < 0.001 |
| | 60 | F5 | 4.482 | < 0.001 |

**Supplementary Table 3.** A summary of the number of contacts of in each region-of-interest, and for each patient. The temporal lobe region-of-interest includes electrodes found in the medial temporal lobe (MTL). Note that the nature of the linear mixed-effects models used in our main analyses means that N is defined by the total number of electrodes in a given region-of-interest rather than the total number of participants with electrodes in a given region.

| Patient | Lobular Region-of-Interest | | | | MTL Region of Interest | | |
|---|---|---|---|---|---|---|---|
| | Frontal | Parietal | Occipital | Temporal | Parahipp. | Hipp. | Amygdala |
| 1 | 51 | 15 | 0 | 3 | 0 | 3 | 0 |
| 2 | 0 | 8 | 16 | 32 | 4 | 4 | 0 |
| 3 | 44 | 12 | 0 | 12 | 0 | 0 | 4 |
| 4 | 28 | 8 | 0 | 28 | 0 | 4 | 4 |
| 5 | 0 | 16 | 0 | 56 | 4 | 4 | 4 |
| 6 | 30 | 3 | 0 | 15 | 0 | 0 | 3 |
| 7 | 40 | 12 | 16 | 28 | 4 | 4 | 0 |
| 8 | 6 | 3 | 0 | 48 | 9 | 6 | 0 |
| 9 | 16 | 0 | 32 | 20 | 4 | 0 | 0 |
| 10 | 24 | 15 | 16 | 21 | 0 | 3 | 0 |
| **Total** | 239 | 92 | 64 | 263 | 25 | 28 | 15 |

**Supplementary Table 4.** Mapping of experiment tasks onto latent variables for experiment 1.

| Experimental task | Latent factor | | |
|---|---|---|---|
| | Head angle | Auditory input | Visual input / eye movements |
| Uncued Head Rotation | ✓ | | ✓ |
| Cued Head Rotation | ✓ | ✓ | ✓ |
| Cued Eye Movements | | ✓ | ✓ |
| Rotation without Visual Input | ✓ | ✓ | |

**Supplementary Table 5.** Mapping of experiment tasks onto latent variables for experiment 2.

| Experimental task | Latent factor | | |
|---|---|---|---|
| | Head angle | Position z-axis | Relocation x-axis |
| Cued Head Rotation | ✓ | | |
| Standing Head Rotation | ✓ | ✓ | |
| Cued Head Rotation at 60° offset | ✓ | | |
| Standing Head Rotation after relocation | ✓ | ✓ | ✓ |

**Supplementary Table 6.** Mapping of training/testing combinations onto latent variables for the dissociation of head rotation and head direction. In short, activity related to head rotation should generalise across sitting positions (i.e., when they faced different angles in the environment); head direction activity should not.

| Training condition | Testing Condition | Latent factor | |
|---|---|---|---|
| | | Head rotation | Head direction |
| Cued Head Rotation | Cued Head Rotation | ✓ | ✓ |
| Cued Head Rotation | Cued Head Rotation at 60° offset | ✓ | |
| Cued Head Rotation at 60° offset | Cued Head Rotation | ✓ | ✓ |
| Cued Head Rotation at 60° offset | Cued Head Rotation at 60° offset | ✓ | |

**Supplementary Table 7.** Mapping of training/testing combinations onto latent variables for the dissociation of location-independent and location-specific head direction signals. In short, activity related to location-independent head direction signals should generalise across conditions, whereas location-specific signals should not.

| Training condition | Testing Condition | Latent factor | |
|---|---|---|---|
| | | Location independent | Location specific |
| Cued Head Rotation | Cued Head Rotation | ✓ | ✓ |
| Cued Head Rotation | Standing Head Rotation | ✓ | |
| Cued Head Rotation | Standing Rot. after relocation | ✓ | |
| Standing Head Rotation | Cued Head Rotation | ✓ | |
| Standing Head Rotation | Standing Head Rotation | ✓ | ✓ |
| Standing Head Rotation | Standing Rot. after relocation | ✓ | |
| Standing Rot. after relocation | Cued Head Rotation | ✓ | |
| Standing Rot. after relocation | Standing Head Rotation | ✓ | |
| Standing Rot. after relocation | Standing Rot. after relocation | ✓ | ✓ |

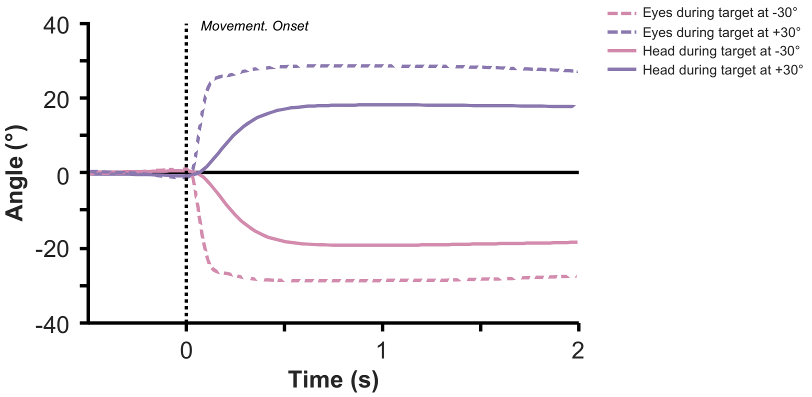

**Supplementary Figure 1. Saccades precede physical rotation during orienting tasks.** Following an auditory cue, participants made saccades/head rotations to the target screen (time = 0 reflects onset of head/ocular movement). The coloured lines depicted mean angular position as a function of time, averaged over trials and participants. Purple lines reflect rotations to the +30° screen; pink lines reflect rotations to the +30° screen. Filled lines reflect head rotations, dotted lines reflect saccades. As the eye-tracker could not track eye position for the far left/far right screen, neither the head direction nor eye position data for these trials are plotted here.

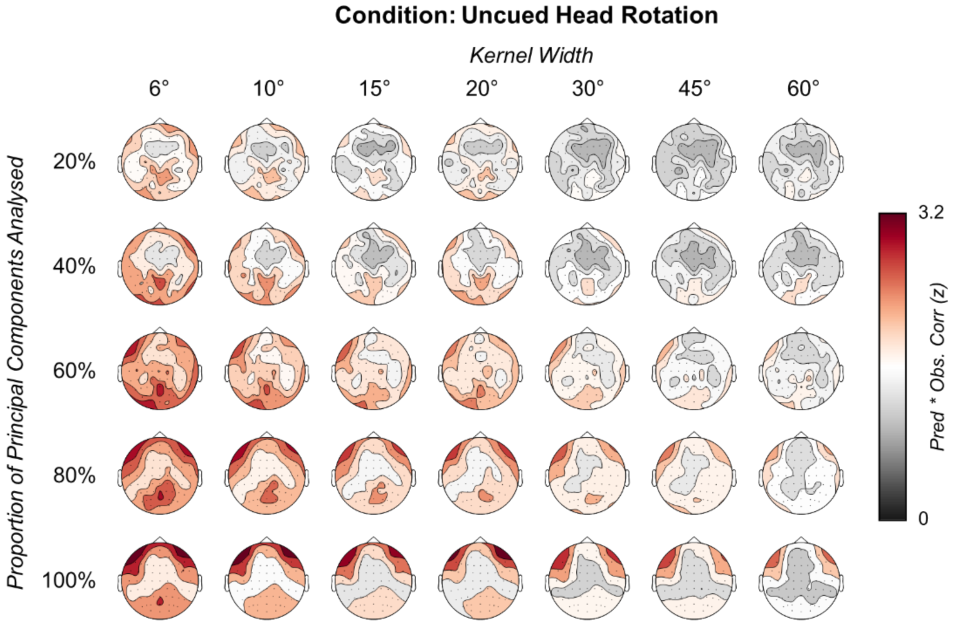

**Supplementary Figure 2. Topographies of the correlation values between FEM-predicted and observed EEG signal for each kernel width and principal component percentage during the "uncued head rotation" task of Experiment 1.** The FEM-predicted signal better correlated with the observed signal when using narrower tuning widths, aligning with the results reported when using the linear mixed-effects models. When using all principal components, a topography reminiscent of saccades could be observed over frontal electrodes, but this declined with the number of EMG-like components removed. In contrast, the peak at Pz remained regardless of the number of EMG-like components removed. These results align with those of the linear mixed-effects model, which found a peak over Pz after regressing out confounds related to muscle activity. For those curious about why the peak effects in these plots are found at 6° and the peak effects for the linear mixed-effects model are found at 20°, this is likely a result of other regressors in the linear mixed-effects models explaining more variance for the narrower tuning widths than the head direction regressor. Indeed, one should interpret these correlation plots with caution as they do not distinguish the effects of head direction from those of visual/saccadic activity, auditory cuing or muscular activity.

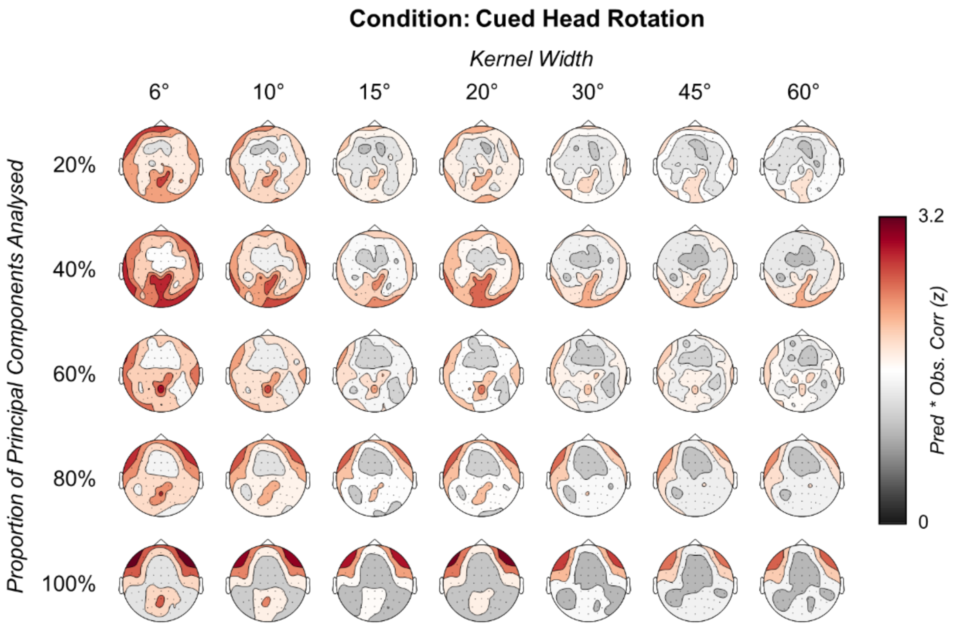

**Supplementary Figure 3. Topographies of the correlation values between FEM-predicted and observed EEG signal for each kernel width and principal component percentage during the "cued head rotation" task of Experiment 1.** For further discussion on this plot, see the legend of supplementary figure 2.

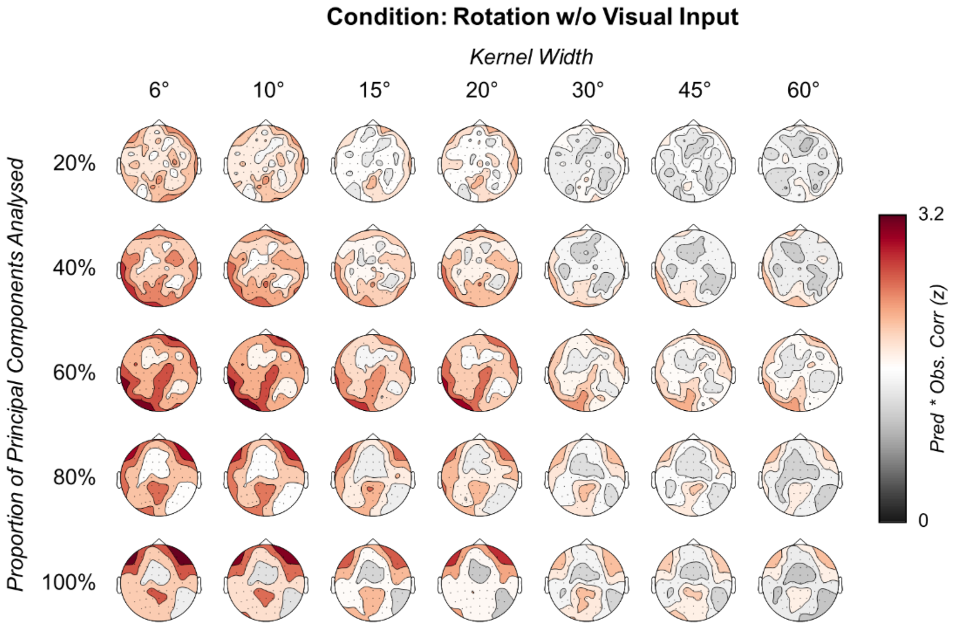

**Supplementary Figure 4. Topographies of the correlation values between FEM-predicted and observed EEG signal for each kernel width and principal component percentage during the "rotation without visual input" task of Experiment 1.** For further discussion on this plot, see the legend of supplementary figure 2.

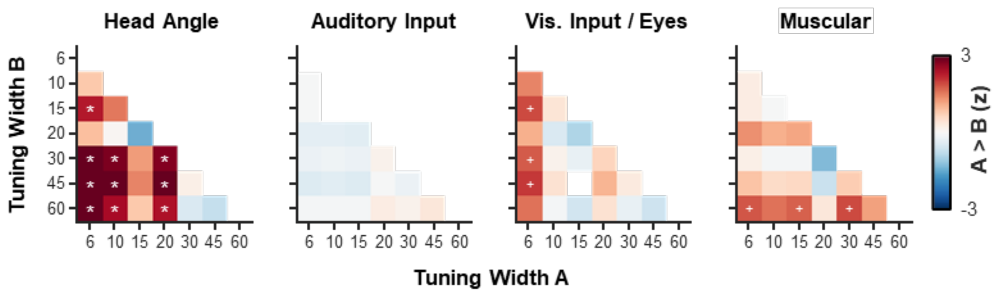

**Supplementary Figure 5. Comparison of FEM performance between tuning widths for all regressors in Experiment 1.** Head angle and visual input/saccade regressors showed a small trend where narrower FEM tunning widths better predicted the EEG data than broader tuning widths. The fact that each model has the potential to explain the same variance may explain why larger effects are not observed. $^{*}p_{FDR} < 0.05$; $^{+}p_{uncorr} < 0.05$.

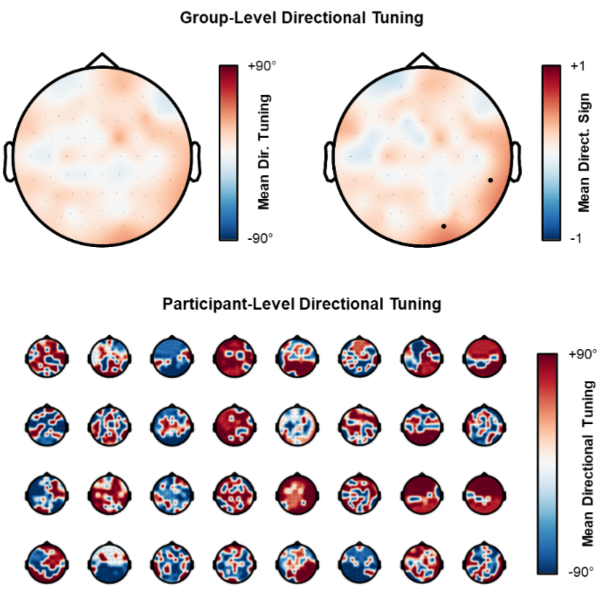

**Supplementary Figure 6. Absence of consistent directional tuning across participants.** Here, the preferred tuning of each electrode was computed for each block and each session and then averaged for each participant individually. The top left plot visualises the mean angle across all participants. The top right plot visualises the mean of the sign of the angle across all participants (+1 indicates that all participants had a tuning angle > 0 for a given electrode; -1 indicates that all participants had a tuning angle < 0 for a given electrode). The black dots indicate channels which showed a sign preference across participants that was above chance ($p_{uncorrected} < 0.05$; no channels survived multiple correction comparison). The participant-level topographies beneath depict the individual preferred tuning angles. A consistent effect across participants is not obvious. The absence of a topographic effect of preferred tuning aligns with earlier rodent work (e.g., Giocomo et al., 2014).

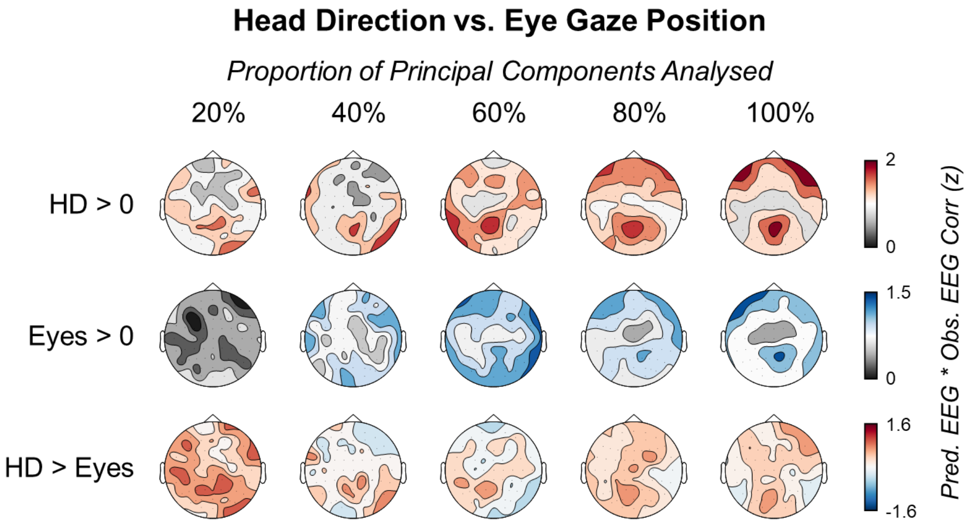

**Supplementary Figure 7. Comparison of FEMs for head direction and eye gaze position.** Two sets of FEMs were created, one which predicted EEG based on current heading angle and the other which predicted EEG based on current eye gaze position. As eyetracking data only existed for trials in which participants saccaded to the -30°/+30° screens, we restricted the head direction FEM to trials in which participants rotated their heads to the same screens to ensure a fair comparison between the models. Despite restricting the head direction FEM to data from the -30°/+30° screens, a significant cluster was observed for all proportions of components (for all PCs: $p_{cluster} < 0.001$), with posterior central electrodes continuing to show the strongest effects. Similarly, the eye gaze-based FEM produced significant clusters in all conditions (for all PCs: $p_{cluster} < 0.001$), though the posterior central dominance was less clear. Critically, a direct contrast of topographies revealed that the head direction FEM outperformed the eye gaze-based FEM for all conditions except when 80% and 20% of components were analysed (100% PCs: $p_{cluster} < 0.001$; 80% PCs: no cluster formed; 60% PCs: $p_{cluster} = 0.002$; 40% PCs: $p_{cluster} = 0.001$; 20% PCs: no cluster formed; though both the 80% and 20% conditions still produced a pattern reminiscent of those seen in the other conditions). Taken together, these results suggest that neural activity is tuned to changes in both heading angle and eye gaze position, but changes in heading angle appears to better predict neural activity than changes in eye gaze position.

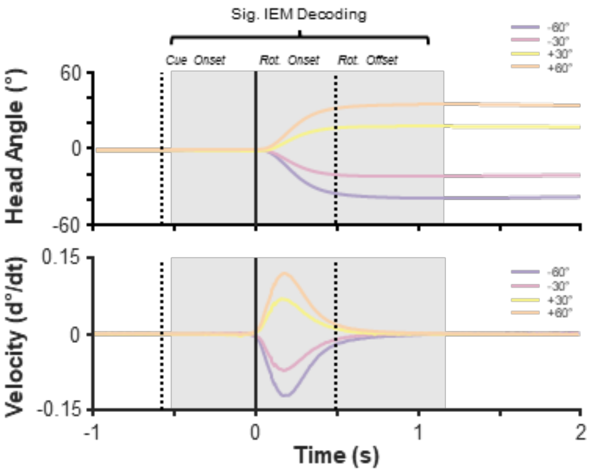

**Supplementary Fig. 8. Time-series of head rotation and rotation velocity.** While head rotation velocity varies between conditions, decodablility of the EEG signal is achievable in windows that extent far beyond when differences in velocity can be observed. This suggests that the encoding model is not simply tracking head velocity. Of course, that is not to say that head velocity does in no way contribute to head direction signals. Indeed, such information may be key to computing head direction (for further details; see Blair and Sharp, 1995).

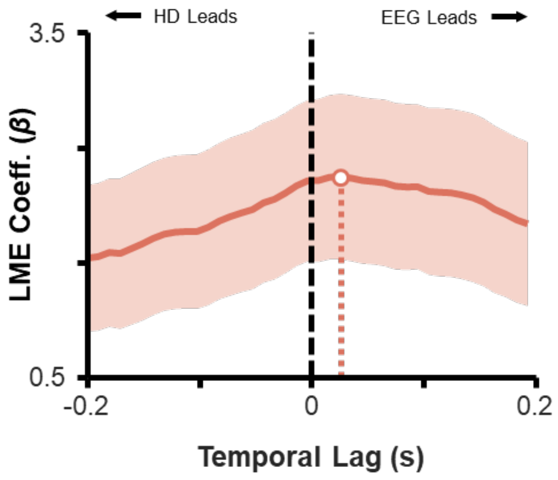

**Supplementary Figure 9. Lag-based analysis of the "uncued head rotation" task in isolation.** Time-series of regressor coefficients derived from linear mixed effects models used to predict lag-based forward encoding model performance. The deep red indicates the estimated regressor coefficients. Error bars indicate the 95% confidence intervals of the coefficients, as computed by the Matlab function fitlme(). Given that the auditory cue was tied to the head rotation from the second task onwards, one could argue that the lag-based effects in the main text were driven by associative memory rather than anticipatory coding. To address this concern, we conducted the lag analysis solely on the "uncued head rotation" task, which had no auditory cuing and occurred before any association between the auditory cue and the head rotation could be formed. In line with the results in the main text, we observed that the EEG leads changes in head direction (max. z = 5.934, $p_{FDR} < 0.001$; lead vs. lag contrast: max. z = 5.183, $p_{FDR} < 0.001$). These results suggest that the lag-based effects cannot be attributed to associative memory between the auditory cue and the head rotation.

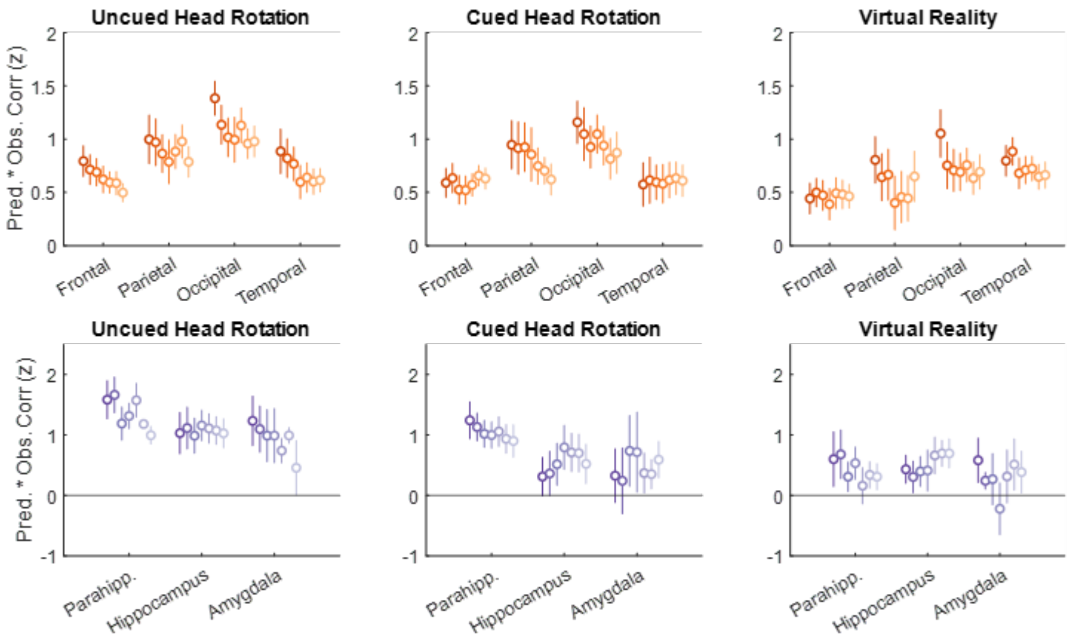

**Supplementary Figure 10. Topographies of the correlation values between FEM-predicted and observed iEEG signal for each kernel width during the "cued head rotation" task.** Error bars depict the standard error of the mean across electrodes. The FEM-predicted signal better correlated with the observed signal when using narrower tuning widths, aligning with the results reported when using the linear mixed-effects models.

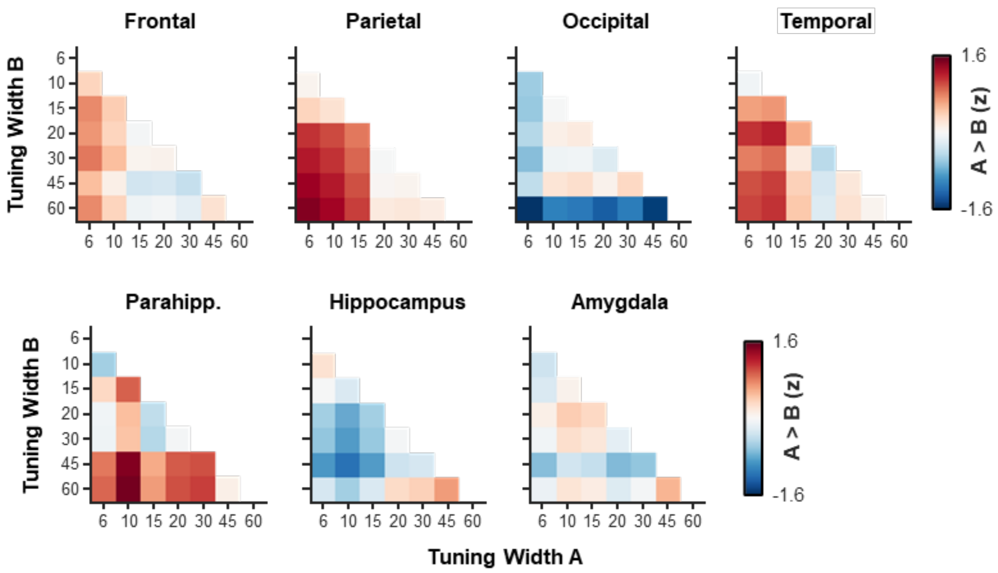

**Supplementary Figure 11. Comparison of FEM performance between tuning widths for all regions-of-interest in the intracranial data of Experiment 1.** While there was a general trend where narrower tuning widths outperformed broader widths, these effects were weak and did not survive multiple comparison correction. The smaller patient sample size (relative to the healthy participants) may mean there is insufficient power detect an effect.

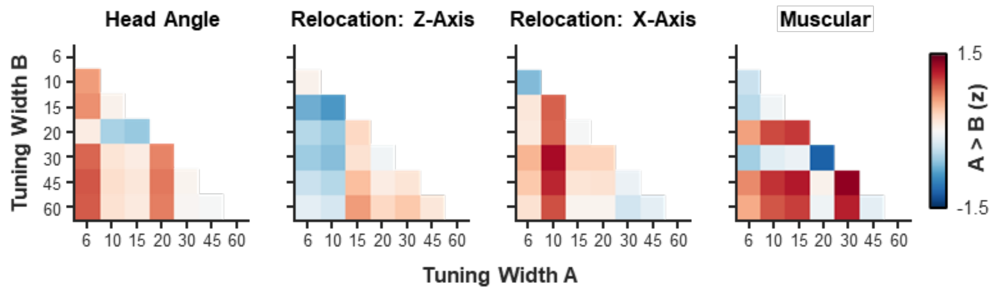

**Supplementary Figure 12. Comparison of FEM performance between tuning widths for all regressors in Experiment 2.** Head angle and visual input/saccade regressors showed a small trend where narrower FEM tunning widths better predicted the EEG data than broader tuning widths. The smaller/differing effects between this experiment and the first may be due to differences in power (Exp. 1: n = 32; Exp. 2: n = 20). The smaller sample size of this experiment was decided upon as it would have sufficient power to replicate the main effect (i.e., FEM performance > 0), not that it would have sufficient power to differentiate FEM performance between different tuning widths (an analysis suggested during the peer review stage).

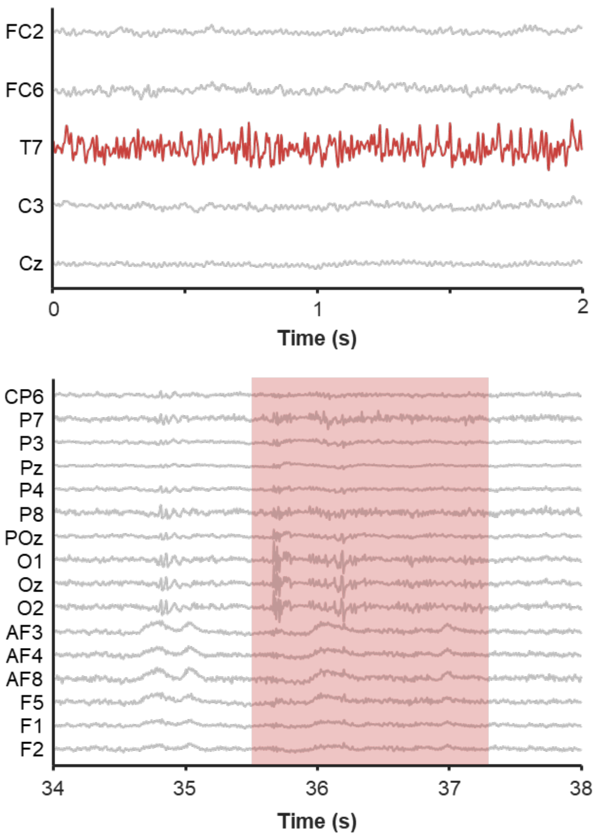

**Supplementary Figure 13. Exemplar EEG artifacts.** In the top panel, a bad channel (red) is visualised in the context of other good channels (grey). Such channel would be interpolated based on the patterns of activity observed in their topographic neighbours. In the bottom panel, a period of artefactual data is visualised (shaded red area). Trials containing such activity would be removed from further analysis.

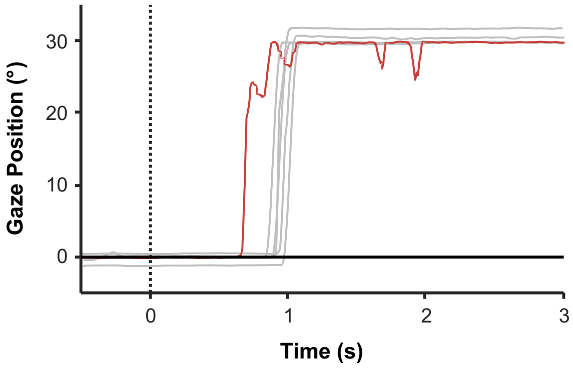

**Supplementary Figure 14. Exemplar eyetracker artifact.** A bad trial (red), consisting of peculiar, physiologically implausible motion, is visualised in the context of other good trials (grey).

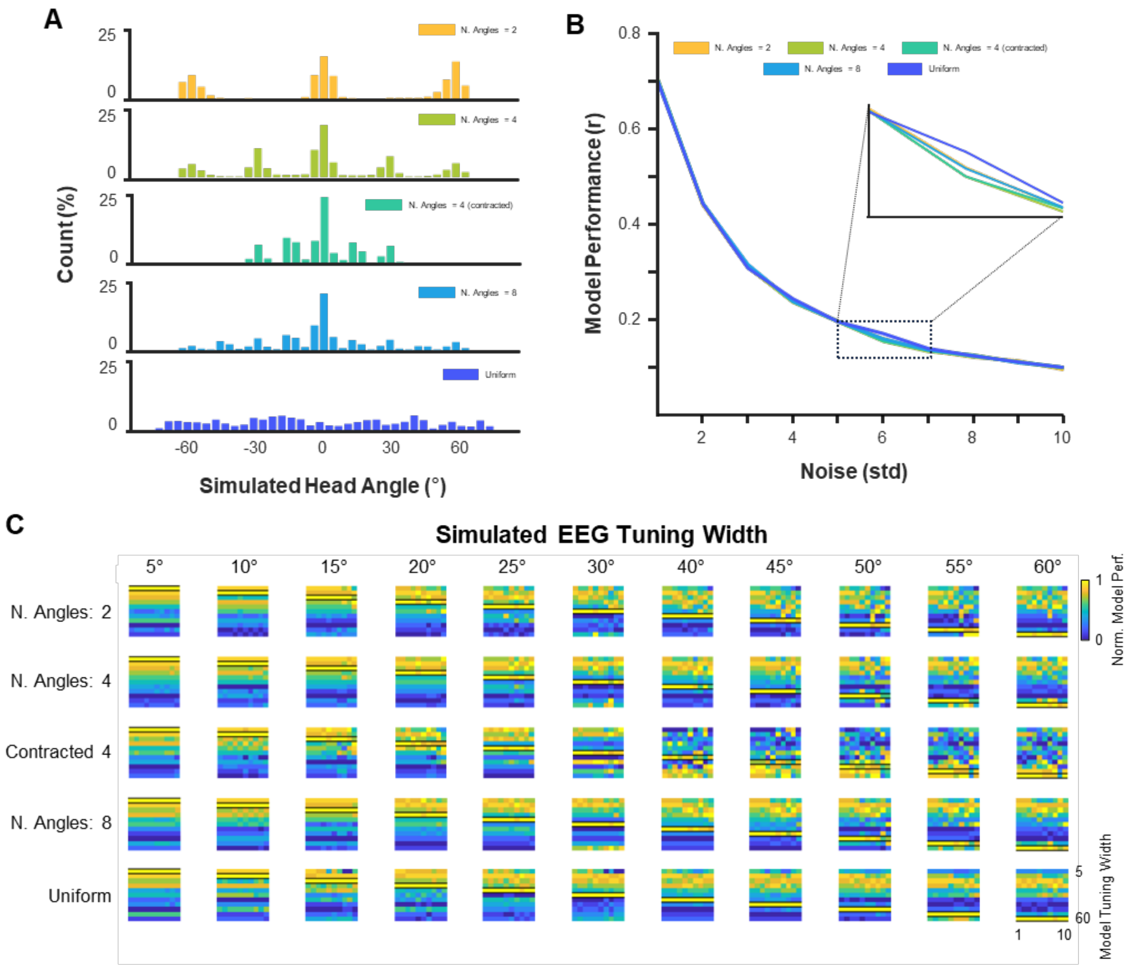

**Supplementary Figure 15. The influence of head direction sampling bias on model performance. (A)** In the main experiment, head direction is not uniformly distributed, but is instead centred on 0°, +30°, +60°, -30°, and -60°. To examine whether this sampling bias influences model performance, we simulated four conditions with differing distributions of head directions. Three conditions took the same form as the main experiment, with set head directions (either two head directions, four head directions [mimicking the main experiment], or eight head directions). In addition to this, we simulated a condition where head directions were contracted (i.e., the angles fell short of the target by ~30%) and another which had an approximately uniform head direction. **(B)** A FEM using 20° kernels was fit to the simulated head direction data in order to predict simulated EEG data that had a tuning width of 20°. Normally-distributed noise was added to the data. As noise increased, model performance decreased for all conditions equally, with the uniform simulation being slightly less susceptible to noise than the other simulations, suggesting that our main experiment was unlikely to overestimate the size of the effect. **(C)** Expanding this analysis to FEMs using a variety of kernel widths (ranging from 5° to 60°) and to simulated EEG with a range of tuning widths (ranging from 5° to 60°) revealed that the FEMs best predicted EEG data when the kernel widths of the model and the tuning widths of the EEG matched, and that this did not vary as a function of bias in the head direction sampling. Altogether, these results suggest that biased sampling of head direction does not impact model performance.

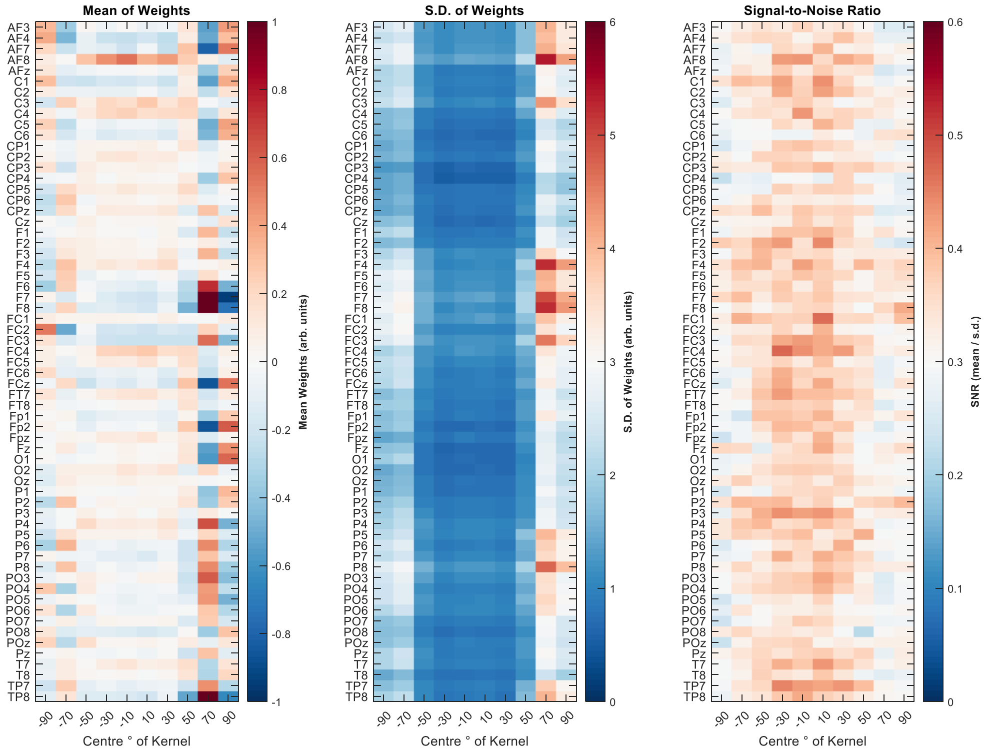

**Supplementary Figure 16. Stability of weights across participants, cross-validation folds and tasks for the scalp EEG data of Experiment 1.** Left plot depicts the mean beta across participants for each channel and each kernel of the 20° FEM model. Middle plot depicts the standard deviation of the beta across participants for each channel and each kernel of the FEM model. Right plot depicts the signal-to-noise ratio (mean divided by standard deviation) for each channel and each kernel of the FEM model. In each plot, the kernel on the far left represents the kernel capturing the most extreme head angles to the left while the kernel on the far right captures the most extreme head angles to the right. The kernels in the centre align with the centre screen. There is a general trend for kernels sitting in the centre of the plot to exhibit less variance and therefore greater signal-to-noise. This is likely because these kernels are well sampled across trials/folds/tasks and therefore produce more robust estimates. Similar results are obtained when looking at the weights from our simulations (see supplementary figure 22).

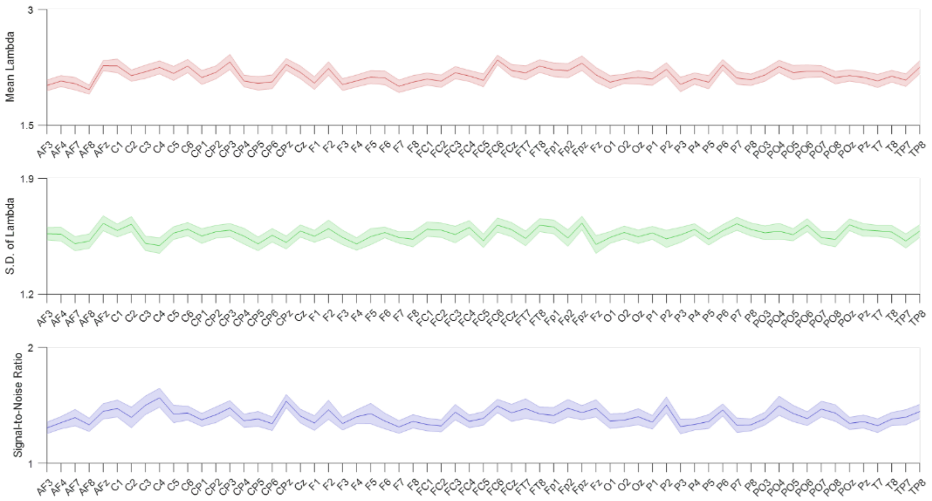

**Supplementary Figure 17. Stability of lambda across participants for the scalp EEG data of Experiment 1.** Each plot depicts the mean (coloured line) and standard error (shaded error) of lambda metrics. The top plot depicts mean lambda across cross-validation runs and tasks, averaged across participants. The middle plot depicts the standard deviation of lambda across cross-validation runs and tasks. The bottom plot depicts the signal-to-noise ratio (mean divided by standard deviation) across cross-validation runs and tasks, averaged across participants.

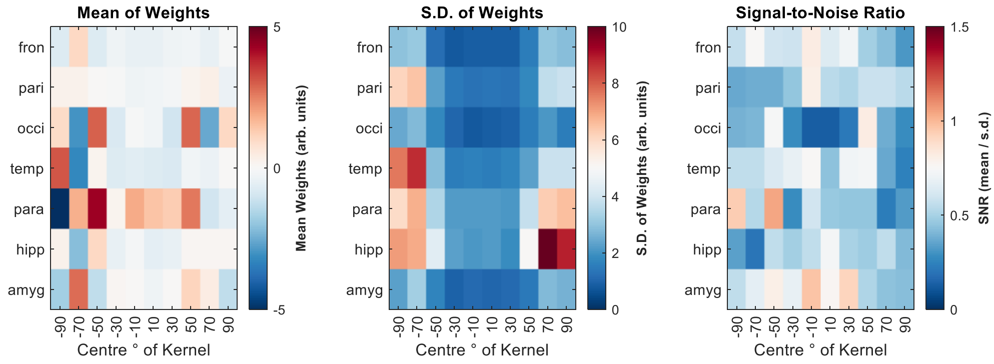

**Supplementary Figure 18. Stability of weights across participants, cross-validation folds and tasks for each region-of-interest of the intracranial EEG data of Experiment 1.** For full legend details, see supplementary figure 16.

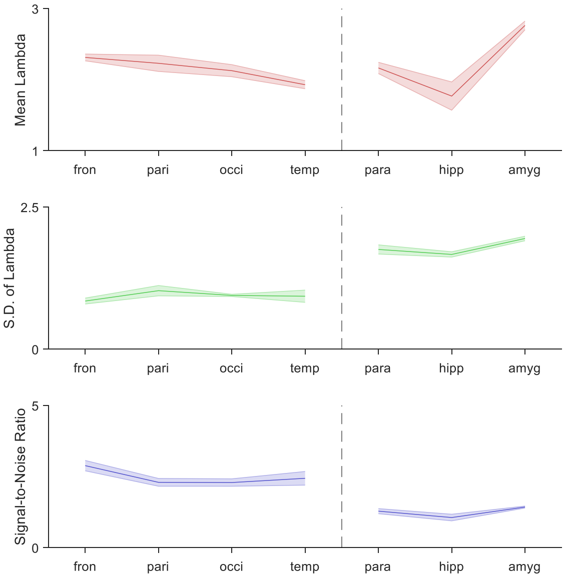

**Supplementary Figure 19. Stability of lambda across participants for each region-of-interest of the iEEG data of Experiment 1.** For full legend details, see supplementary figure 17.

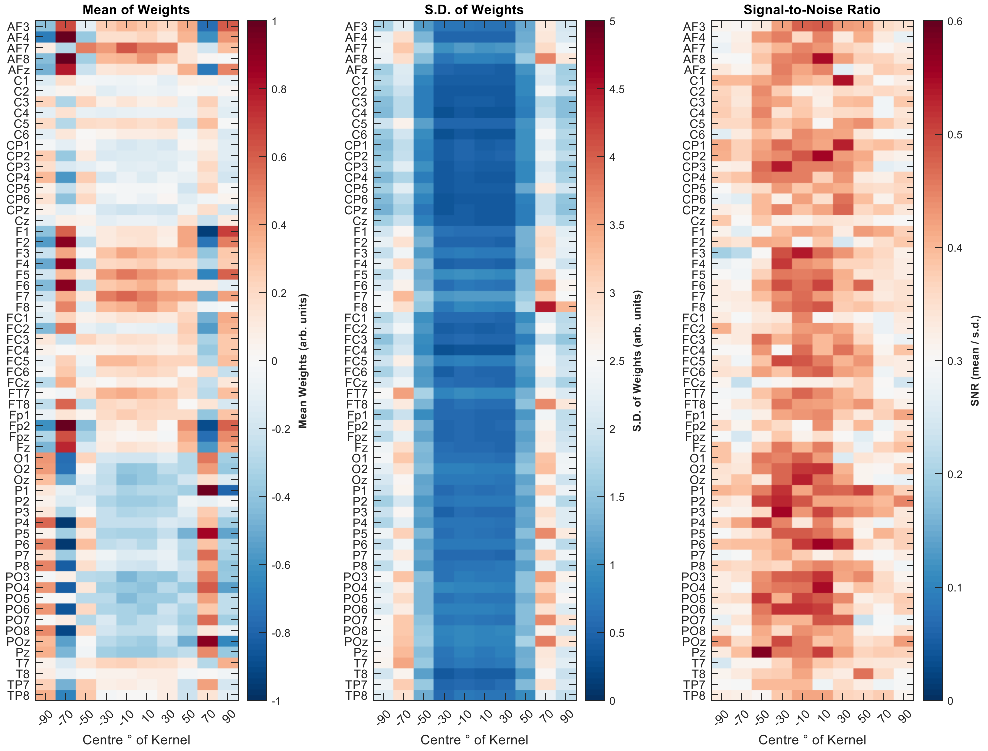

**Supplementary Figure 20. Stability of weights across participants, cross-validation folds and tasks for the scalp EEG data of Experiment 2.** For full legend details, see supplementary figure 16.

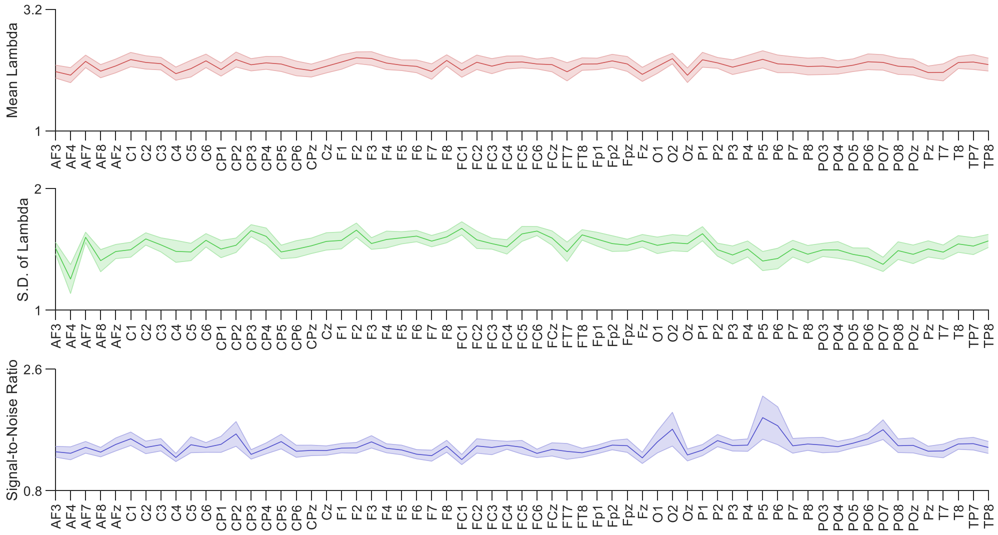

**Supplementary Figure 21. Stability of lambda across participants for each channel of the scalp EEG data of Experiment 2.** For full legend details, see supplementary figure 17.

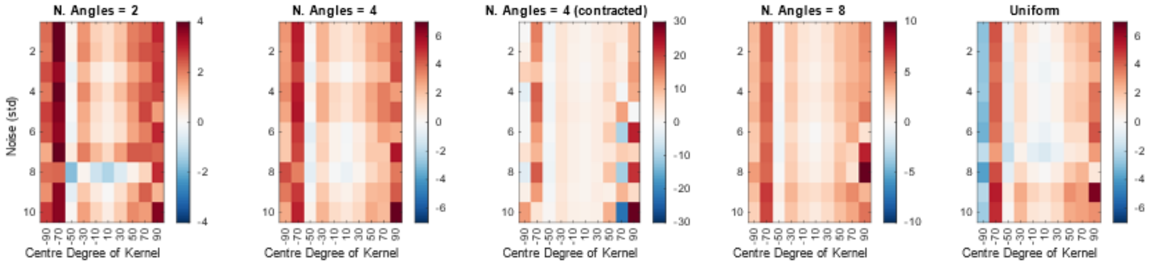

**Supplementary Figure 22. Weight values for simulated models.** Each simulation demonstrates a tendency to heavily up-weight or down-weight the outer kernels where sampling is sparse. Despite this, model performance is consistent across distributions (see supp. figure 15). This suggests that the model is not skewed by outlying or extreme head angles.
